# *Enterocloster citroniae* and related gut microbiome species modulate *Vibrio cholerae* biofilm formation through the production of bioactive small molecules

**DOI:** 10.64898/2025.12.15.694346

**Authors:** Heidi Pauer, Saeideh Nasiri, Nathalia Santos Magalhães, Vivian T. Nguyen, Nicole Victor Ferreira, Larissa D. Silva Ferreira, Athena Brianne Bradshaw, Katherine E. Kirby, Tyler Sabapathy, Chukwuma Great Udensi, Viktoriia Feofanova, Daniel Andrade Moreira, Thiago Estevam Parente, Jacob Wilde, David T. Pride, Emma Allen-Vercoe, L. Caetano M. Antunes

**Author notes:** These authors contributed equally to this work (Heidi Pauer, Saeideh Nasiri).

## Abstract

Cholera is a diarrheal disease that affects millions of people globally. Although the causative agent, *Vibrio cholerae*, has been extensively studied in isolation, investigation of its interactions with the gut microbiota started relatively recently. We and others previously showed that microbiota-derived metabolites significantly influence *V. cholerae* behavior. By investigating how an organic extract of human feces affects *V. cholerae* gene expression, we showed that gut metabolites strongly suppress swimming motility, a trait important for host colonization. Interestingly, extracts of pure cultures of a gut commensal, *Enterocloster citroniae*, recapitulated this inhibition. Here, we present a comprehensive examination of the effect of small molecules produced by *E. citroniae* and related species on *V. cholerae* behavior. We show that *E. citroniae* small molecules inhibit motility by various *V. cholerae* strains, and that several phylogenetically related species produce this activity, although the magnitude of the effect varies between strains. Using biofilm formation assays in static and flow conditions, we show that *V. cholerae* strongly induces biofilm formation in response to *E. citroniae* metabolites. Transcriptome and reporter analyses showed that several genes involved in synthesis of an extracellular polysaccharide are induced by *E. citroniae* metabolites. Lastly, we show that *V. cholerae* interactions with host cells are also modulated by this commensal. These findings advance our understanding of microbiome-pathogen interactions and how commensal bacteria influence *V. cholerae* virulence through the production of small molecules. In the future, this knowledge may be used to design novel microbiome-based therapeutic approaches to combat cholera and other infections.

**Importance:** The human gut is home to a dense and rich community of microbes termed microbiota. This community has critical functions for host health, including protection against enteric pathogens. Despite this important role, we have only recently scratched the surface of the interactions that occur between members of the microbiota and pathogenic invaders. Cholerae is a disease that still causes significant morbidity and mortality worldwide. Studying how the causative agent, *Vibrio cholerae*, interacts with the microbiota will have implications not only for our understanding of this important microbial community, but may also lead to the development of new therapeutic strategies against cholera and potentially other infectious diseases.

## Introduction

*Vibrio cholerae* is the causative agent of cholera, a diarrheal disease that continues to cause significant morbidity and mortality worldwide (1). The behavior of *V. cholerae*, including how it employs multiple virulence factors to cause disease, has been extensively studied in pure cultures. Recently, studies have begun to explore *V. cholerae* interactions with host-associated microbes, but our understanding remains incomplete (2–5). The gut microbiota, a complex community of microbes residing in the gastrointestinal tract of all animals, plays a vital role in protecting the host from infection, including by shaping host immune responses and by directly inhibiting pathogen growth and virulence (6). Among the many ways in which the microbiota affects infection, one prominent mechanism is through the production of bioactive compounds that influence pathogen virulence and colonization (7–9). Recent studies have described the role of microbiota-derived metabolites in influencing the virulence of *V. cholerae*. *Bacteroides vulgatus* has been shown to restrict *V. cholerae* growth during infection of antibiotic-treated mice through the production of short-chain fatty acids (3). *Blautia obeum*, on the other hand, inhibits *V. cholerae* colonization by inhibiting virulence factor gene expression through the production of a quorum sensing signal (2). However, interactions between microbiota members and *V. cholerae* are not limited to those that inhibit pathogen virulence. For instance, interactions between *V. cholerae* and *Paracoccus aminovorans* have been shown to enhance biofilm formation, which facilitates colonization and persistence in the host (10). Therefore, complex interactions between *V. cholerae* and microbiome members likely occur during gut colonization.

To investigate the impact of microbiome-derived small molecules on *V. cholerae* behavior, we previously determined the impact of an organic extract of human feces on *V. cholerae* global gene expression (8). By doing so, we found that the fecal metabolome is a strong repressor of *V. cholerae* motility, a key virulence factor that facilitates host colonization and transmission of *V. cholerae* (11). Prior to this report, our group had described that *Enterocloster citroniae*, a member of the human gut microbiome, produces small molecules that inhibit invasion gene expression by *Salmonella enterica* (7). Therefore, we tested whether the *E. citroniae* strain with activity against *S. enterica* was also capable of recapitulating the effect of the human fecal extract on *V. cholerae* motility. Indeed, organic extracts of *E. citroniae* cultures strongly inhibit *V. cholerae* swimming motility (8). In this study, we describe a comprehensive examination of the effect of small molecules produced by *E. citroniae* on various aspects of *V. cholerae* behavior. We examine the effects of *E. citroniae* extracts on *V. cholerae* global gene expression, biofilm formation, toxin production, and interactions with host cells. We also describe the effect of several gut commensals phylogenetically related to *E. citroniae* on *V. cholerae*. Altogether, our results expand our knowledge of the effects microbiome members impinge upon various aspects of *V. cholerae* virulence through the production of small molecules. Studying these phenomena will generate new insights into *V. cholerae* interactions with the microbiome and may uncover new targets for the development of microbiome-related therapeutics.

## Materials and methods

### Bacterial strains and growth conditions

Strains used in this study are listed in **Table S1**. *V. cholerae* was routinely grown in Luria-Bertani (LB; Fisher Bioreagents, Waltham, MA, USA) or Brain Heart Infusion (BHI; MIDSCI, Fenton, MO, USA) medium, as indicated. BHI broth used for *V. cholerae* growth was supplemented with 0.5% yeast extract (MIDSCI) and hemin (5 µg/mL; Sigma-Aldrich, St. Louis, MO, USA). Cultures were incubated at 37°C with shaking at 200 rpm. *Enterocloster* spp. and *Clostridium innocuum* strains were cultured in BHI supplemented with 0.5% yeast extract, cysteine (1 g/L; Thermo Scientific, Waltham, MA USA), hemin (5 µg/mL), and menadione (0.1 mg/mL; Sigma-Aldrich) at 37°C in an anaerobic chamber (Coy Laboratory Products, Ann Arbor, MI, USA) under an anaerobic atmosphere composed of 80% N_2_, 5% H_2_, 15% CO_2_.

For growth curve assays, an overnight culture of *V. cholerae* in BHI was subcultured 1:200 into 200 µL of BHI broth containing either an extract from one of the commensal strains studied (*Enterocloster* spp., *C. innocuum*), or a control extract, prepared as described below, in a 96-well polystyrene plate (TPP, Trasadingen, Switzerland). Cultures were incubated at 37°C and bacterial growth was monitored by measuring the optical density at 600 nm (OD_600_) using a SpectraMax i3 spectrophotometer (Molecular Devices, San Jose, USA) for 24 hours with readings taken every 30 minutes.

### Isolation of gut microbes

Strains were isolated from donor stool samples by dilution and culture on fastidious anaerobe agar (LabM, Potters Bar, England) containing 5% defibrinated sheep’s blood (Hemostat Laboratories, Dixon, USA), and incubation at 37°C for up to 72 hours under an atmosphere of CO_2_:H_2_:N_2_ 5:5:90 in a Baker Ruskinn Concept 500 anaerobic chamber (Baker, Sanford, USA). Isolated colonies were purified by repeated plate subculture under the same growth conditions and subsequently stored at −80°C. Genomic DNA was extracted from overnight plate cultured cell biomass using a Maxwell® 16 DNA purification kit (Promega) and identified by sequencing of the amplified 16S rRNA gene using Sanger sequencing (as a service through the University of Guelph Advanced Analysis Centre). Species identifications were determined using NCBI BLAST (www.ncbi.nlm.nih.gov/BLAST/).

### Chemicals

Short-chain fatty acids (SCFAs; Thermo Scientific) were prepared as concentrated stock solutions (1 M) of sodium acetate (99% pure), sodium propionate (99% pure), and sodium butyrate (≥98% pure) in ultrapure water, filter-sterilized (0.22 µm), and stored at −20°C until used. Stocks were diluted directly into BHI medium immediately before use to achieve final concentrations that varied based on the SCFA (acetate: 15, 30, 60 mM; butyrate: 5, 10, 20 mM; propionate: 5, 10, 20 mM), or mixed at the ratios specified in the text and figure legends. Antibiotics were obtained from Sigma-Aldrich and used at the following concentrations: kanamycin, 100 µg/mL; chloramphenicol, 10 µg/mL; polymyxin B, 20 µg/mL.

### Plasmids and DNA manipulation

All plasmids used in this study are listed in **Table S1**. *Escherichia coli* S17-λpir harboring pCMW5 (kindly provided by Dr. Christopher M. Waters), which confers constitutive expression of the green fluorescent protein (GFP), was used as the donor strain to mobilize the plasmid into *V. cholerae* C6706 via conjugation. The *vpsR* reporter plasmid was constructed using pBBRLux (12). The upstream sequence of *vpsR* (−495 to the start codon) was cloned into the *Bam*HI and *Spe*I restriction sites of pBBRLux by GenScript (Piscataway, NJ, USA), to generate pLC010 (P*_vpsR_-luxCDABE*). The plasmid was first transformed into *E. coli* BW29427 and then transferred into *V. cholerae* C6706 by conjugation, as follows. All strains were grown overnight to the stationary phase at 37°C with shaking (200 rpm). *V. cholerae* C6706 was subcultured (1:100 dilution) and grown in LB medium at 37°C with shaking until reaching an OD_600_ of approximately 0.7. *E. coli* and *V. cholerae* cells were washed with LB medium by centrifugation (5 min at 13,000 *x*g) to remove antibiotics and were concentrated 5-fold and 20-fold in LB medium, respectively. Fifteen microliters of each culture were mixed and spotted onto LB agar plates without antibiotics. When using the BW29427 strain, an auxotrophic *E. coli* strain requiring 2,6-diaminopimelic acid (DAP), 300 µg/mL of DAP was added to the plates. Conjugation mixtures were incubated overnight at 37°C. The bacterial spots were scraped from the agar and resuspended in 1 mL of LB medium. One hundred microliters of the suspension were plated onto agar plates supplemented with the appropriate antibiotic corresponding to the resistance marker in the respective plasmid and incubated overnight at 37°C. Colonies were picked the day after, grown overnight, and stored at −80°C.

### Small molecule extraction

Small molecule extraction was performed as described previously (13). *Enterocloster* and *C. innocuum* strains were inoculated onto anaerobic blood agar plates (Remel, Lenexa, KS, USA) and incubated at 37°C under anaerobic conditions for 48 hours. A single isolated colony was then inoculated into a small volume of BHI broth (∼5 mL) and incubated for 24 hours at 37°C under anaerobiosis. Subsequently, a subculture was prepared by diluting this culture 1:50 into a larger volume of BHI and incubating it anaerobically at 37°C for an additional 24 hours. Cultures of *Enterocloster* and *C. innocuum* were then extracted with ethyl acetate (Fisher Chemical, Waltham, MA). To do this, ethyl acetate was added to the culture at a 1:1 (v/v) ratio. The mixture was shaken vigorously and allowed to settle for 10 minutes; this process was then repeated once. The organic phase was then collected and transferred to a clean bottle. Control extracts were produced by extracting sterile culture media using the same procedure. Extracts were stored at −20°C for up to 4 weeks. Before experiments, the solvent was evaporated using an R-300 rotary evaporator (BÜCHI, Flawil, Switzerland) or a Savant SPD140DDA SpeedVac concentrator (Thermo Fisher Scientific, Waltham, MA, USA) at 40°C. Dried residues were resuspended in culture media, and the pH was adjusted to 7.2 before sterilization through 0.2-μm filters. The relative concentrations of extracts were defined based on the volume of organic phase recovered after extraction. Extracts resuspended in a volume of culture medium equal to the recovered organic phase were regarded as 1X. Bacterial culture and control extracts were used at a 2X relative concentration in all experiments.

### Motility assays

To observe *V. cholerae* swimming motility, BHI plates containing 0.3% agar were prepared containing either control extract or extracts from *Enterocloster* or *C. innocuum* strains. The medium was poured into 100-mm Petri dishes and allowed to solidify overnight at room temperature. Plates were gently inoculated with a single colony picked from a plate containing fresh, overnight growth using a sterile toothpick and incubated at 37°C for 18 hours. Swimming was quantified by measuring the diameter of bacterial growth (8). The effect of SCFA on *V. cholerae* motility was assessed under the same conditions.

For the microscopic observation of swimming motility, a GFP-labeled *V. cholerae* strain (VC002; C6706 with pCMW5) was grown in BHI broth containing either a control extract or an extract from *E. citroniae* WAL17108 for 3 hours at 37°C with shaking (200 rpm). After incubation, the cultures were diluted to an OD_600_ of 0.8. A 1-µL aliquot of the culture was placed on a microscope slide, covered with a coverslip, and visualized using a Leica DM 5500 B microscope with a 60x objective (Leica Microsystems, Deerfield, USA). Videos were then recorded for 10 seconds. Four biological replicates were conducted, and, for each replicate, four fields were recorded. Bacterial swimming was analyzed using TrackMate, an ImageJ Fiji plugin (14).

### Biofilm formation assay

Biofilm formation assays (crystal violet staining) were conducted as described previously (13), with some modifications. Briefly, *V. cholerae* C6706 was grown overnight in LB broth at 37°C with shaking (200 rpm). Following overnight growth, the culture was diluted 1:200 in BHI broth containing either a control extract or an extract from *E. citroniae* WAL17108 and grown as static cultures in a 96-well polystyrene plate (TPP) at 37°C for 24 hours. After incubation, planktonic bacteria were removed from the microplate by aspiration using a micropipette, and the wells were washed three times with phosphate-buffered saline (PBS pH 7.4; Fisher Bioreagents). The biofilm was fixed at 60°C for 1 hour and stained with 200 µL of a 0.1% crystal violet (Sigma-Aldrich) solution for 15 minutes at room temperature. Excess stain was removed by rinsing the wells three times with PBS. The dye was then solubilized using 200 µL of a 95% ethanol solution, and the OD_570_ was measured in a Varioskan LUX Multimode Microplate Reader (Thermo Fisher Scientific).

Biofilm formation was also analyzed under flow conditions using the microfluidic BioFlux 200 system (Cell Microsystems, Durham, NC). *V. cholerae* VC002 was grown overnight in LB broth at 37°C with shaking (200 rpm). Following overnight growth, the culture was diluted 1:100 in BHI broth containing either a control extract or an extract from *E. citroniae* WAL17108, and this bacterial suspension was used as the inoculum. Prior to inoculation, the channels of a 48-well Low Shear Plate (Cell Microsystems) were primed with 50 μL of PBS to fill the channels and remove air from the system. After priming, 20 µL of bacterial inoculum was added to the outlet well and perfused from the outlet to the inlet at 1 dyn/cm² until cells filled the viewing windows. The plate was incubated statically for 1 hour at 37°C to allow for microbial attachment. One milliliter of BHI broth containing either a control extract or an extract from *E. citroniae* WAL17108 was then added to the inlet wells of each channel. To remove non-adherent cells, the channels were perfused from the inlet to the outlet at 1 dyn/cm² for 1 minute. Then, the shear was reduced to 0.5 dyn/cm^2^ and channels were perfused with medium for 18 hours. To monitor biofilm formation, the plate was mounted on the stage of an Olympus IX81 Inverted Microscope (Evident, Waltham, USA). Biofilm formation was tracked over 18 hours, with images collected every hour using a 10x objective at two individual positions in multiple channels. After image acquisition, a representative region was selected per channel, and the average pixel intensity (mean gray value) was measured for each sample using ImageJ. The Corrected Total Cell Fluorescence (CTCF) was calculated using the following formula: CTCF = integrated density - (area of selected cell × mean fluorescence of background readings).

### RNA sequencing and data analysis

*V. cholerae* C6706 was grown overnight in LB medium at 37°C with shaking (200 rpm). The culture was diluted 1:200 in BHI broth containing either a control extract or an extract from *E. citroniae* WAL17108 and incubated at 37°C with shaking (200 rpm). After approximately 3 hours of growth (mid-log phase), bacterial RNA was stabilized using RNAprotect Bacteria Reagent (Qiagen, Germantown, MD, USA) according to the manufacturer’s instructions. Bacterial cells were pelleted by centrifugation, and total RNA was isolated using the SV Total RNA Isolation System (Promega, Madison, WI, USA), following the manufacturer’s recommendations. RNA samples were treated with Turbo DNase (2 U/μL; Life Technologies, Waltham, MA, USA) to eliminate residual DNA, followed by an additional purification step using the SV Total RNA Isolation System (Promega), according to the manufacturer’s recommendations. RNA concentration and purity were assessed using an ND-1000 NanoDrop spectrophotometer (Thermo Fisher Scientific). Absence of DNA contamination in the RNA was checked by quantitative Real-Time PCR, as described in the next subsection. RNA sequencing (RNA-seq) was performed on an Illumina NextSeq 2000 platform using a P1 flow cell, as single reads (100 bp), at The University of Kansas Genome Sequencing Core. After sequencing, the quality of reads was assessed using FastQC v0.12.1 (http://www.bioinformatics.babraham.ac.uk/projects/fastqc/) and MultiQC v1.14 (15). Reads were then mapped against the *V. cholerae* C6706 reference genome (GCF_015482825.1) using STAR software v2.7.10b (parameter: --quantMode GeneCounts) (16). Raw counts were normalized and the differentially expressed genes were determined with DESeq2 R package (Log2 fold-change ≥1, p-adjusted <0.05) (17). Raw data was deposited in Gene Expression Omnibus (https://www.ncbi.nlm.nih.gov/geo/), under study accession number GSE313194.

### Quantitative Real-Time PCR (qRT-PCR)

*V. cholerae* C6706 cultures were prepared under the same conditions described for RNA-seq. Bacterial RNA was stabilized using RNAprotect Bacteria Reagent (Qiagen) and extracted with the SV Total RNA Isolation System (Promega), according to the manufacturer’s protocols. Following DNase treatment and RNA cleanup (as described above), cDNA synthesis was performed using Go Script^TM^ Reverse Transcription System (Promega, Madison, WI, USA). qRT-PCR was conducted on a QuantStudio™ 3 Real-Time PCR System (Applied Biosystems, Waltham, MA, USA) using PowerTrack™ SYBR™ Green Master Mix (Applied Biosystems, Waltham, MA, USA). The absence of DNA contamination in RNA samples was checked by performing cDNA synthesis reactions without reverse transcriptase in parallel to the ones containing enzyme and then checking for the absence of amplification during qRT-PCR. Gene expression levels were normalized to the housekeeping gene *recA*, and relative expression was calculated using the 2^-ΔΔCt^ method (18). Primer sequences are listed in **Table S1**.

### Transcriptional reporter assays

Luminescence assays were performed using *V. cholerae* C6706 carrying the pLC010 *vpsR* transcriptional reporter plasmid (designated *V. cholerae* VC007). Overnight cultures were grown in LB medium supplemented with chloramphenicol (10 µg/mL) at 37°C with shaking (200 rpm). Cultures were diluted 1:200 into fresh BHI medium with chloramphenicol (10 µg/mL and containing either a control extract or an extract from *E. citroniae* WAL17108. Cultures were incubated at 37°C in white, 96-well plates (BRAND GmbH & Co KG, Wertheim, Germany) with intermittent shaking (5 seconds right before each reading). Luminescence and OD_600_ were measured every 30 minutes for 16 hours using a SpectraMax i3 plate reader (Molecular Devices). Luminescence is expressed as relative light units (RLU), calculated by dividing the luminescence signal by the corresponding OD_600_ to normalize for cell growth.

### Cell culture

Human colonic epithelial cells (HT-29) were obtained from the American Type Culture Collection (ATCC; Manassas, VA, USA), and were routinely cultured in Dulbecco’s Modified Eagle Medium (DMEM; Gibco, Thermo Fisher Scientific) supplemented with 10% (v/v) heat-inactivated fetal bovine serum (FBS; Cytiva, Marlborough, MA, USA), 1% (v/v) GlutaMAX (Gibco), 1% (v/v) non-essential amino acids (Gibco), and 1% (v/v) penicillin (100 U/mL) and streptomycin (100 μg/mL) (Gibco). Cells were maintained at 37°C in a humidified atmosphere containing 5% CO_2_ until reaching approximately 80-90% confluence.

### Cell viability and cytotoxicity assays

Cell viability and cytotoxicity were evaluated using 3-(4,5-dimethylthiazol-2-yl)-2,5-diphenyltetrazolium bromide (MTT) and lactate dehydrogenase (LDH) release assays following exposure of HT-29 cells to control extract or extracts of *E. citroniae* WAL17108. After 22-24 h of treatment under standard culture conditions (37°C, 5% CO₂), cell viability was determined by the MTT colorimetric assay. Briefly, cells were incubated with an MTT solution (0.5 mg/mL; Sigma-Aldrich, St. Louis, MO, USA) for 2 hours. The medium was then removed and the resulting formazan crystals were solubilized in DMSO. Absorbance was measured at 570 nm using a Varioskan LUX microplate reader. Vell viability (%) was calculated as: cell viability (%) = (absorbance of treated cells × 100)/ absorbance of untreated cells. Cytotoxicity was quantified by measuring LDH release into the culture supernatant using the CytoTox 96^®^ Non-Radioactive Cytotoxicity Assay (Promega, Madison, WI, USA) according to the manufacturer’s instructions. Absorbance was measured at 490 nm. LDH release (%) was calculated relative to unexposed cells (0% cytotoxicity) and cells treated with a lysis solution (100% cytotoxicity). Increased LDH release was interpreted as an indicator of membrane damage.

### Adhesion and invasion assays

*V. cholerae* adhesion and invasion assays were performed using HT-29 cells seeded in 24-well plates at 10^6^ cells/well in supplemented DMEM. *V. cholerae* strains were grown overnight in LB broth at 37°C with shaking (200 rpm). Overnight cultures were subcultured 1:200 into BHI broth, containing either an extract from *E. citroniae* WAL17108 or a control extract, and grown at 37°C with shaking (200 rpm) to the mid-log growth phase (OD_600_ of approximately 0.5). Cultures were washed once with sterile PBS and resuspended in supplemented DMEM (no antibiotics). HT-29 monolayers were infected at a multiplicity of infection (MOI) of 200:1 and incubated for 1 hour at 37°C. For adhesion assays, following infection, monolayers were washed once with PBS to remove non-adherent bacteria. Cells were lysed with 0.025% Triton X-100 (Sigma-Aldrich) in PBS for 5 minutes at room temperature. Lysates were serially diluted and plated on LB agar to enumerate adherent bacteria by determining the number of colony forming units (CFUs). For invasion assays, after the 1-hour infection period, cells were washed once with PBS and incubated for an additional hour in DMEM containing gentamicin (100 µg/mL) to kill extracellular bacteria. Cells were then washed with PBS and lysed with 0.025% Triton X-100 for 5 min. Intracellular bacteria were quantified by plating serial dilutions of the lysates on LB agar.

### Bacterial intracellular survival assay

Bacterial survival assays were performed to evaluate the persistence of *V. cholerae* C6706 within HT-29 cells over time in the presence or absence of *E. citroniae* WAL17108 extracts or a control extract. HT-29 cells were seeded into 24-well plates (10^6^ cells/well) and grown for 18 hours to reach 80-90% confluence. Cells were maintained at 37°C in supplemented DMEM in a 5% CO_2_ atmosphere. *V. cholerae* cultures were prepared as described for adhesion and invasion assays. Bacteria were washed once with PBS and resuspended in antibiotic-free DMEM, containing either an extract from *E. citroniae* WAL17108 or a control extract. HT-29 monolayers were infected at a MOI of 200:1 in the continued presence or the respective extracts. After a 1-hour infection period at 37°C, monolayers were washed once with PBS to remove non-adherent bacteria. Fresh supplemented DMEM containing gentamicin (100 µg/mL) and the appropriate extract (control or *E. citroniae*) were added, and plates were incubated further. At 2 hours post-infection, cells were washed once with PBS and the medium was replaced with supplemented DMEM containing gentamicin (10 µg/mL) and either a control or *E. citroniae* extract. The infections continued until 6 hours. Monolayers were then washed once with PBS and lysed with 0.025% Triton X-100 for 5 minutes, and lysates were serially diluted and plated for CFU enumeration as described above.

### Statistical analyses

Statistical significance was determined using unpaired *t* tests or multiple unpaired t tests. For multiple comparisons, analysis of variance (ANOVA) was used with Bonferroni’s, Dunnett’s, or Geisser-Greenhouse correction, as indicated. Mann Whitney tests and Welch’s t tests were performed for biofilm, motility, adhesion, invasion, and reporter assays, as indicated. Statistical analyses were performed using Prism 10 (GraphPad Software). Specific details of statistical tests, statistical significance values (“p”), sample sizes (‘‘n’’), and replicates are indicated in figure legends. For all analysis, significance was considered as *p*<0.05.

### Ethics statement

Stool samples from which strains ET129A-DCM99, ET325A-NB6, ET129A-LIQR2, ET144A-DCM5, ET233A-FMU6 and ET325-BBE13 were collected in a study approved by the University of California San Diego Administrative Panel on Human Subjects in Medical Research under protocol #160524. This study did not require informed consent on behalf of the study subjects because it involved the use of existing records and specimens that were recorded in such a manner that the subjects could not be identified directly or through identifiers linked to the subjects.

## Results

### Bioactive molecules from gut commensals inhibit *V. cholerae* motility

We previously showed that a strain of *E. citroniae*, FM-V5-E, produces compounds that inhibit *V. cholerae* swimming motility (8). For these original studies, we used VC833, a multidrug-resistant *V. cholerae* strain isolated from human feces during a cholera outbreak in Nigeria (19). To determine whether this is a strain-specific phenomenon, we assessed the ability of *E. citroniae* extracts to modulate swimming motility of other *V. cholerae* strains, include 1 Classical and 3 El Tor biotype strains (**Table S1**). Swimming assays were performed on 0.3% BHI agar plates containing either a control extract or an extract from *E. citroniae* FM-V5-E. Exposure to the *E. citroniae* extract significantly reduced the motility diameter of all *V. cholerae* strains tested compared with the control (**Figure 1**). Next, to determine whether this effect is specific to *E. citroniae* FM-V5-E or prevalent among other *E. citroniae* strains as well as other genetically related species of gut commensals, extracts from additional *E. citroniae* strains, strains of other *Enterocloster* species (*E. aldenensis*, *E. bolteae*, *E. clostridioformis*), and strains of *C. innocuum* were tested against *V. cholerae* C6706. As can be seen in **Figure 2**, with the exception of *E. clostridioformis*, all species had at least one strain that displayed activity against *V. cholerae* swimming. However, the prevalence of bioactivity varied between species; while all *E. citroniae* (5) and *E. aldenensis* (5) strains showed activity, most *E. bolteae* (6/9) and all *E. clostridioformis* (2) strains were inactive. Interestingly, *C. innocuum* showed high variation, with some highly active strains and some completely inactive ones. In total, among the 32 strains examined, 20 significantly decreased *V. cholerae* motility (**Figure 2**).

**Figure 1.**
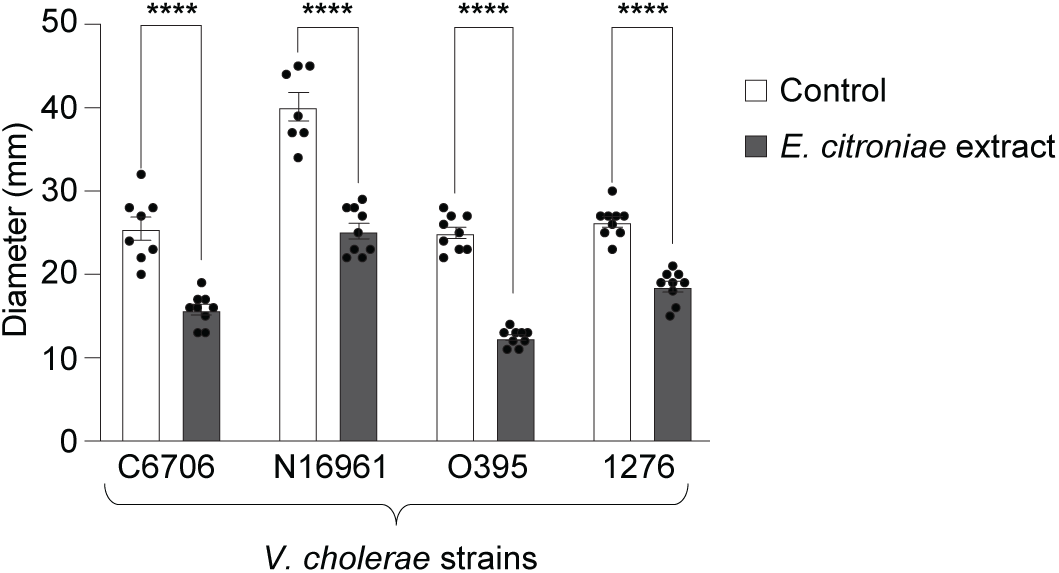
Bioactive molecules produced by *E. citroniae* inhibit swimming motility of multiple *V. cholerae* strains. Swimming motility of *V. cholerae* strains C6706, N16961, O395, and 1276 were assessed on 0.3% agar BHI plates containing either a control extract or an extract from *E. citroniae* FM-V5-E. The diameter of motility zones was measured after 18 hours of incubation at 37°C. Each data point represents an independent biological replicate (*n*=9). Bars indicate the mean ± SEM. Statistical significance was determined by an unpaired *t*-test; \*\*\*\**p*<0.0001 compared with the corresponding control.

**Figure 2.**
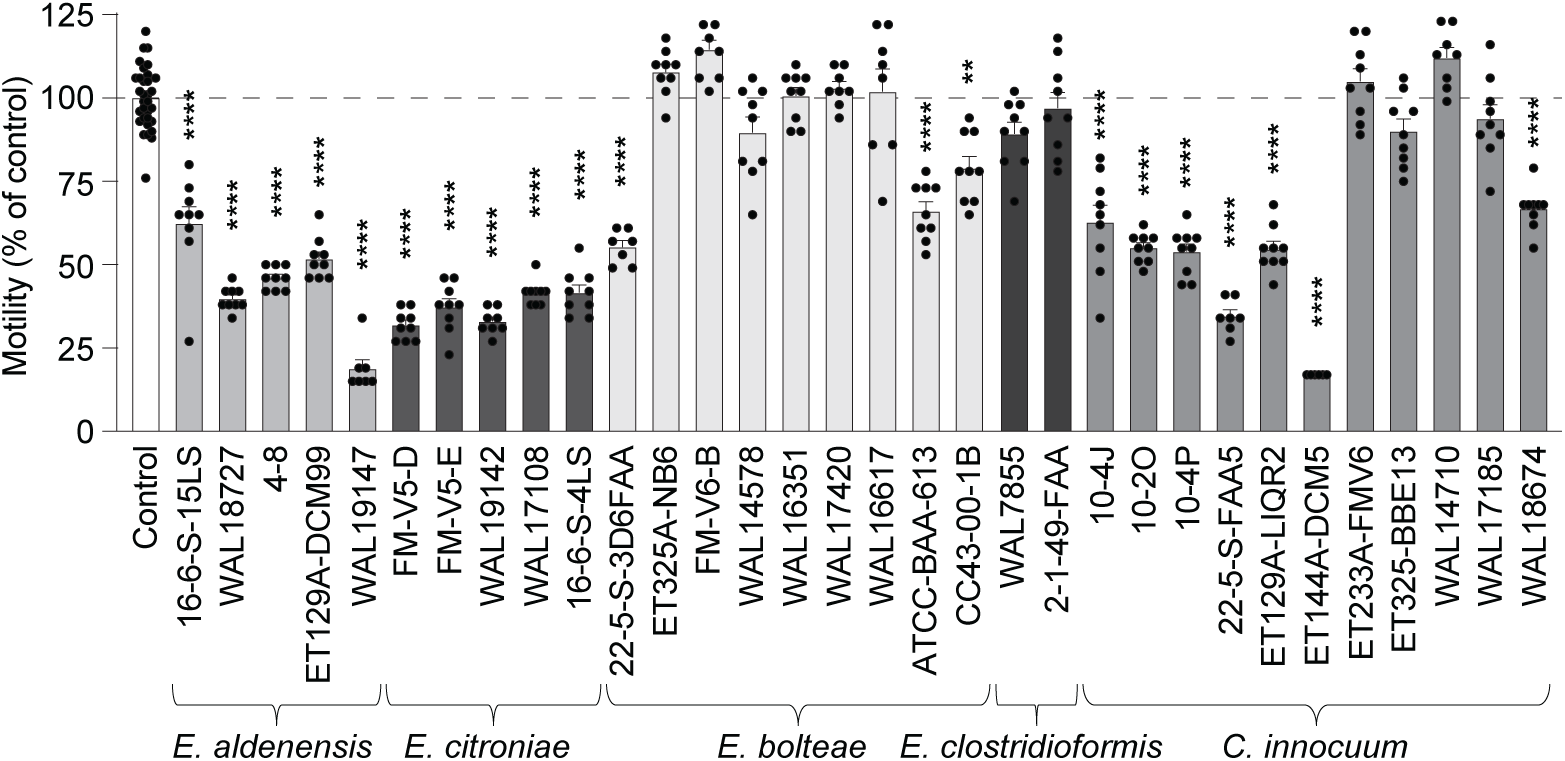
Multiple bacterial species phylogenetically related to *E. citroniae* inhibit *V. cholerae* motility. Swimming motility of *V. cholerae* C6706 is expressed as a percentage of control after exposure to extracts from 32 gut isolates, including *E. aldenensis*, *E. citroniae*, *E. bolteae*, *E. clostridioformis*, and *C. innocuum*. Bars show mean ± SEM. Statistical significance was determined by one-way ANOVA with Dunnett’s multiple-comparisons test; \*\**p*<0.005, \*\*\*\**p*<0.0001. Each data point represents an independent biological replicate (*n*=9).

To directly observe the effect of microbiome-derived bioactive compounds on *V. cholerae* motility, at the cellular level, we tracked fluorescently labeled *V. cholerae* C6706 (VC002) in the presence or absence of a control extract or extracts from *E. citroniae* WAL17108 [**Figure 3; Movies S1 (control) and S2 (*E. citroniae* extract)**]. Consistent with the results obtained from motility assays in agar plates, *V. cholerae* grown in the presence of *E. citroniae* extracts exhibited reduced swimming distance (**Figure 3A**). Additionally, increased bacterial aggregation was observed in cultures treated with *E. citroniae* extracts (**Figure 3B**) compared to the control, suggesting that biofilm formation may also have been affected.

**Figure 3.**
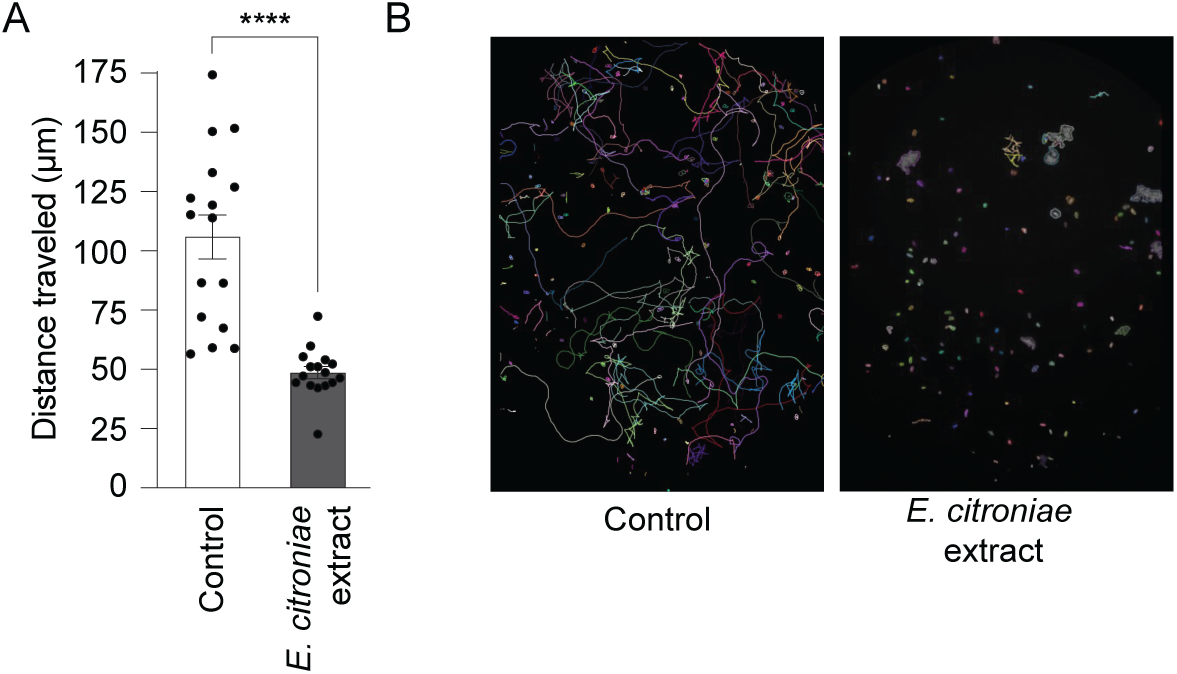
Bioactive molecules produced by *E. citroniae* reduce swimming distance and promote aggregation of *V. cholerae*. (A) Quantification of swimming distances of fluorescently labeled *V. cholerae* VC002 tracked by live imaging in the presence or absence of *E. citroniae* WAL17108 extract or a control extract. Data are mean ± SEM; ****p < 0.0001 (Welch’s t test). Four biological replicates were conducted, and for each replicate, four fields were recorded. (B) Representative motility tracks and fluorescence micrographs showing reduced motility and increased aggregation upon exposure to the extract.

### *E. citroniae* produces bioactive molecules that increase biofilm formation

To determine whether *E. citroniae* metabolites influence *V. cholerae* biofilm formation, we performed a crystal violet staining assay in polystyrene, 96-well plates. Indeed, *V. cholerae* C6706 grown in the presence of an *E. citroniae* extract (WAL17108) displayed significantly higher biofilm biomass compared with the control (**Figure 4A**). To further characterize this phenotype, biofilm formation was analyzed in real time using the BioFlux microfluidic system, which allows dynamic visualization of biofilm development under controlled shear flow. *V. cholerae* VC002 successfully formed biofilms under a shear stress of 0.5 dyne/cm^2^ in both conditions; however, fluorescence quantification after 18 hours revealed a substantial increase in biofilm accumulation in the presence of *E. citroniae* extracts (**Figure 4B**). Representative fluorescence images confirm the enhanced surface coverage of *V. cholerae* biofilms when exposed to the *E. citroniae* extract (**Figure 4C**). Time-lapse movies of GFP-labeled biofilms can be seen in **Movie S3**.

**Figure 4.**
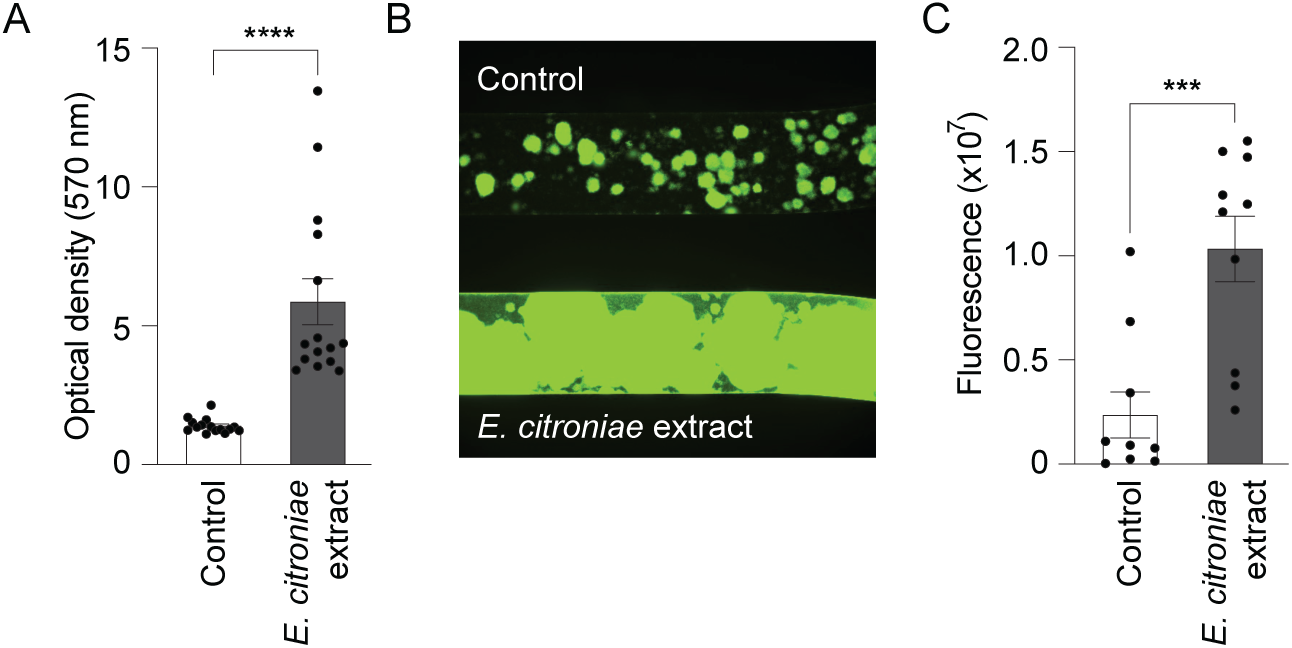
Bioactive molecules produced by *E. citroniae* enhance biofilm formation by *V. cholerae*. (A) Biofilm biomass was quantified by crystal violet staining after 18 h of static incubation at 37 °C in BHI medium containing either an extract from *E. citroniae* or a control extract. Optical density was measured at 570 nm; ****p < 0.0001 (Mann–Whitney test). Three biological replicates were performed, each consisting of five technical replicates. (B) Quantification of fluorescence intensity from GFP-labeled *V. cholerae* VC002 biofilms formed in the BioFlux system after 18 h under 0.5 dyne/cm² flow; ***p < 0.001 (Mann–Whitney test). Five biological replicates were performed, with two images collected per replicate. (C) Representative fluorescence images showing increased biofilm density in the presence of *E. citroniae* extract.

### Microbiome-derived extracts do not markedly affect *V. cholerae* growth

To determine whether gut microbiota members produce compounds that significantly affect *V. cholerae* growth, bacterial cultures were incubated with or without extracts from *Enterocloster* spp. or *C. innocuum* strains, and *V. cholerae* growth was assessed through measurements of the OD_600_ of cultures overtime, in a 96-well plate using a plate reader. As shown in **Figure 5**, *V. cholerae* growth was only slightly delayed in the presence of extracts from *E. citroniae* WAL17108. Other *E. citroniae* and *E. aldenensis* strains also only slightly delayed *V. cholerae* growth (**Figure S1A**), particularly during the exponential phase of growth. In contrast, extracts from *E. clostridiformes*, *E. boltae*, and *C. innocuum* did not affect bacterial growth (**Figures S1B and S1C**). These results suggest that the effects observed on motility and biofilm formation are not due to general effects on bacterial growth.

**Figure 5.**
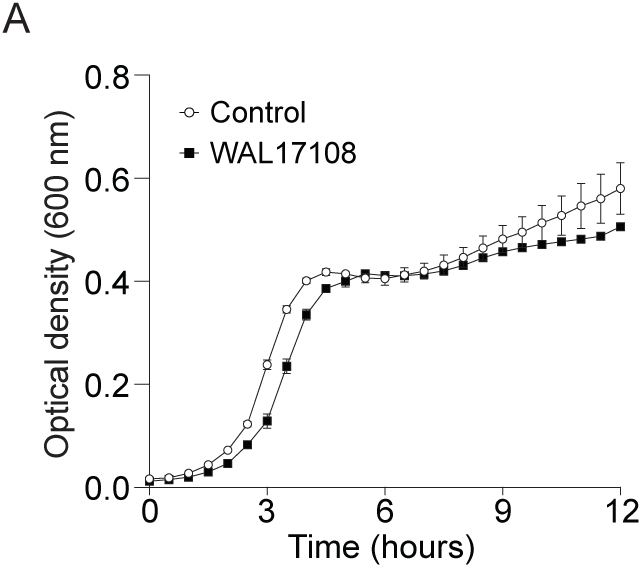
*E. citroniae* extracts do not significantly impact *V. cholerae* growth. Cultures were grown in BHI broth containing either an extract from *E. citroniae* WAL17108 or a control extract at 37°C, and growth was measured by absorbance (OD_600_) for 12 h in a microplate reader. Results represent the average of three independent cultures, each consisting of two technical replicates (n=6), and bars show the standard error of means.

### Short-chain fatty acids do not account for the bioactivity of *E. citroniae* extracts

Many members of the gut microbiome are known to produce short-chain fatty acids (SCFA), which have been extensively shown to display bioactivity [reviewed in (20)]. To assess whether the phenotypic effects of *E. citroniae* extracts on *V. cholerae* could be attributed to SCFAs, we tested the effects of acetate, propionate, and butyrate, individually and in combination, across a range of physiologically relevant concentrations, on *V. cholerae* swimming motility. As shown in **Figure 6A**, none of the tested SCFAs significantly altered *V. cholerae* motility compared with the control condition. Similarly, biofilm formation was not enhanced by SCFA treatment (**Figure 6B**), in contrast to the marked induction observed in the presence of *E. citroniae* extracts. Also, growth curve analyses showed butyrate and propionate, as well as SCFA combinations that included these two SCFA, caused marked reductions in *V. cholerae* growth (**Figure 6C**). These findings indicate that SCFAs alone are not responsible for the inhibition of swimming motility and promotion of biofilm formation by *V. cholerae* caused by *E. citroniae* extracts, and that other bioactive metabolites must be responsible for modulating *V. cholerae* behavior.

**Figure 6.**
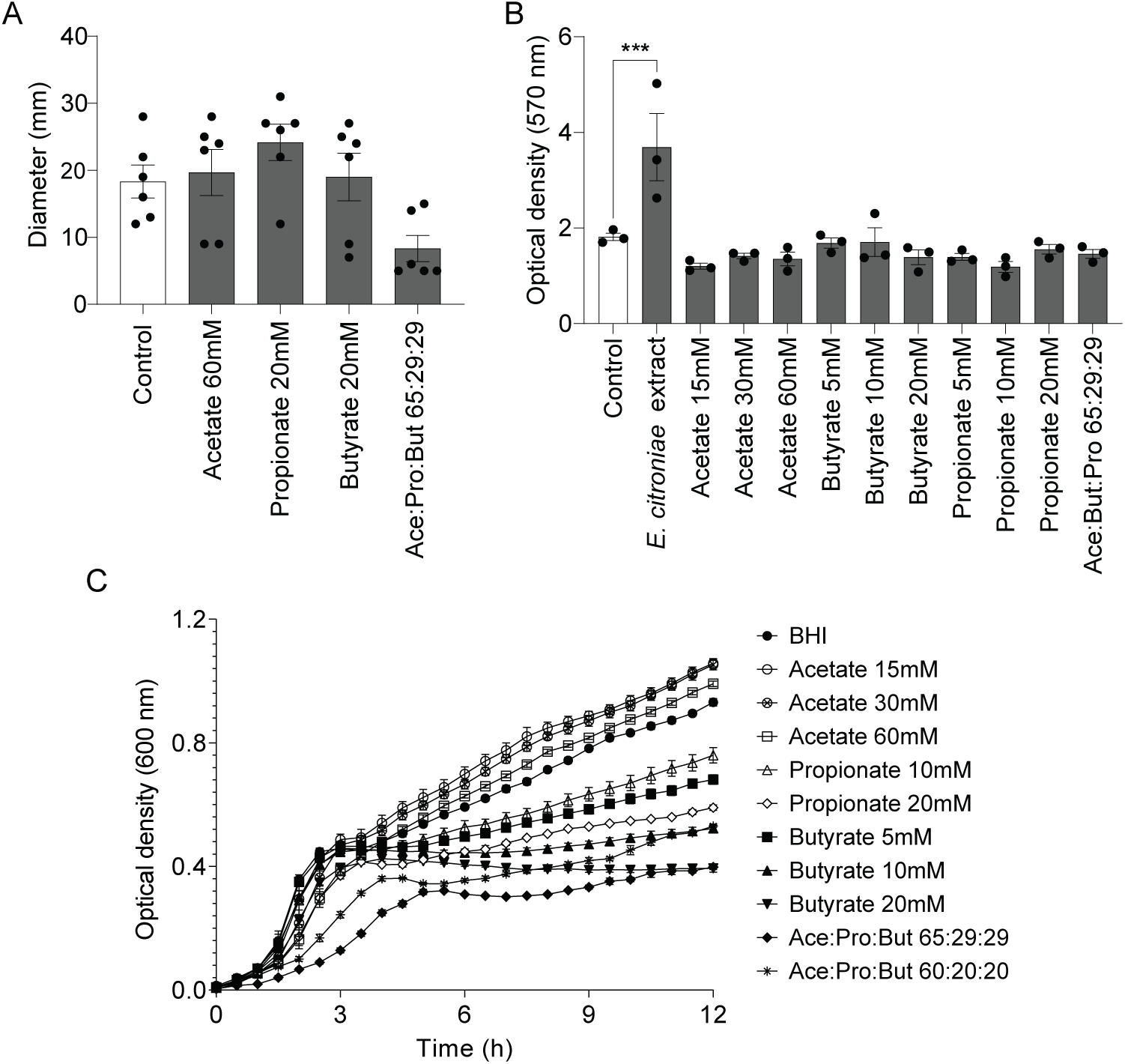
Short-chain fatty acids are not responsible for the effects of *E. citroniae* extracts on *V. cholerae* motility and biofilm formation. (A) Swimming motility of *V. cholerae* strain C6706 was assessed on 0.3% BHI agar plates supplemented with SCFAs at the indicated concentrations. The diameter of motility zones was measured after 18 h of incubation at 37 °C. Each data point represents an independent biological replicate (*n* = 6). Bars indicate the mean ± SEM. One-way ANOVA determined statistical significance; (B) Quantification of biofilm biomass by crystal violet staining (OD_570_ nm) after 18 h of static growth at 37 °C in BHI supplemented with SCFAs at the indicated concentrations. ***p < 0.001 compared with control (One-way ANOVA). (C) Growth curves of *V. cholerae* in BHI supplemented with SCFAs at the indicated concentrations, monitored by OD₆₀₀ over 12 h.

### *E. citroniae* molecules modulate *V. cholerae* global gene expression

To assess the impact of *E. citroniae* metabolites on *V. cholerae* transcriptional activity, we compared the transcriptome of cells grown to the mid-log growth phase in the presence of an *E. citroniae* extract (WAL17108) versus a control extract. Sequencing produced approximately 126 million high-quality reads, with a unique mapping rate over 92.7% to the reference genome (**Table S2**). A total of 576 genes (approximately 15% of the genome) were differentially expressed in response to the *E. citroniae* extract, including 238 upregulated and 338 downregulated genes (**Table S2**). Functional categorization of differentially expressed genes using EggNOG (21) revealed that, among upregulated genes, the most affected categories were energy production and conversion (13.4%), amino acid transport and metabolism (12.4%), and cell wall/membrane/envelope biogenesis (10.6%) **(Figure 7)**. In contrast, downregulated genes were enriched for functions related to carbohydrate transport and metabolism (8.7%), nucleotide transport and metabolism (5.6%), replication/recombination/repair (5.6%), and cell motility (4%) **(Figure 7)**. Notably, genes associated with biofilm formation and nitrate reductase activity were upregulated. Conversely, several virulence-associated *loci* - including components of the type VI secretion system (T6SS), RTX toxin, and toxin-coregulated pilus (*tcp*) cluster - were significantly downregulated, indicating that *E. citroniae* metabolites suppress key virulence pathways in *V. cholerae* **(Table 1)**. To confirm the transcriptomics results, relative mRNA levels of selected genes were quantified by qRT-PCR during *V. cholerae* growth in the absence or presence of an *E. citroniae* extract. The extract significantly upregulated genes associated with biofilm formation (*vpsR*, *vpsT*, *vpsL*, *rbmA*, *bap1*), whereas virulence-related genes, including *tcpA* and *rtxC*, were markedly downregulated **(Figure 8)**. These findings are consistent with the RNA-seq data, further supporting that *E. citroniae*-derived metabolites repress virulence pathways while promoting biofilm-associated gene expression in *V. cholerae*.

**Figure 7.**
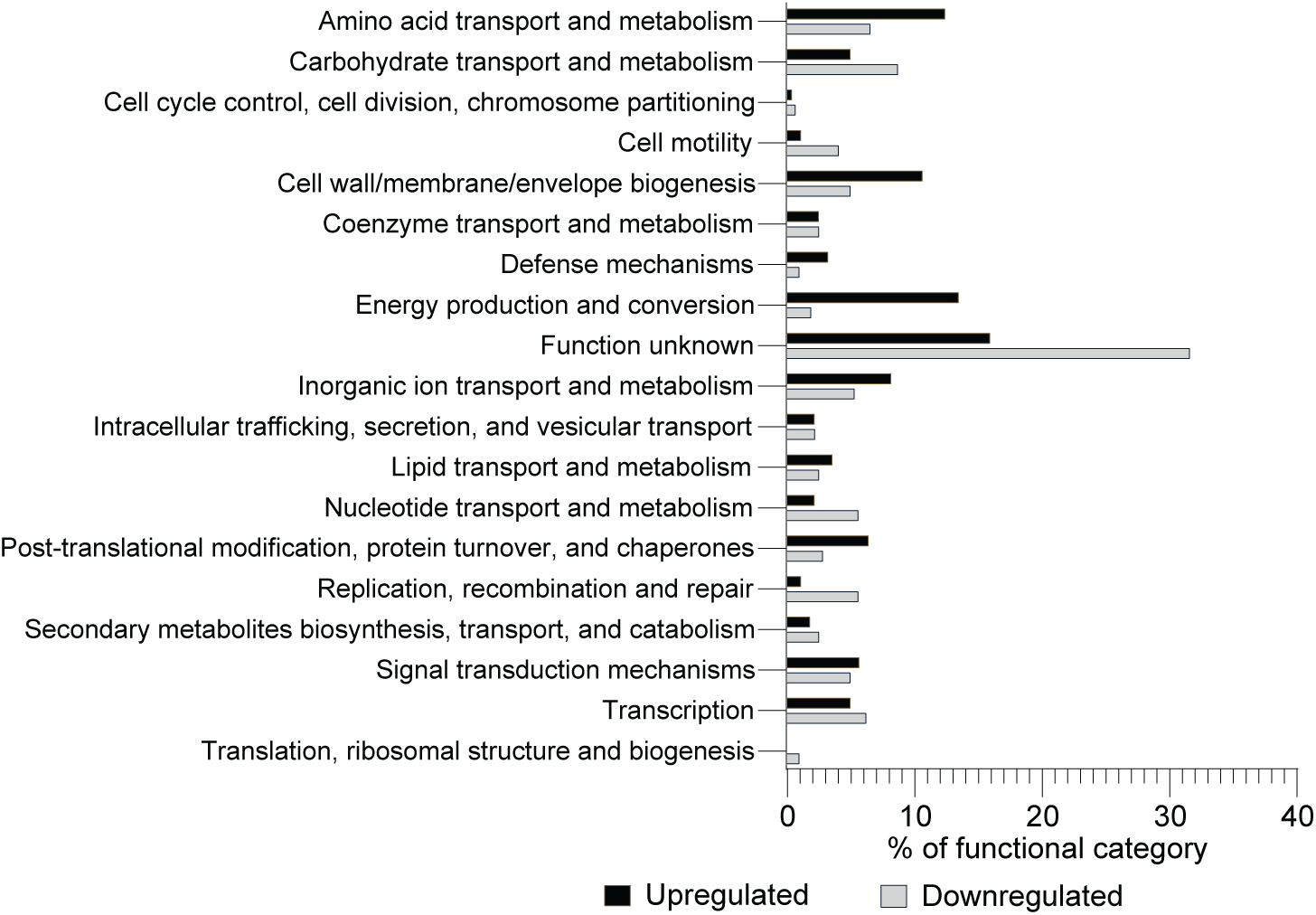
Functional categorization of *V. cholerae* genes differentially expressed during growth in the presence of *E. citroniae* extracts. Genes showing ≥2-fold changes in expression (based on RNA-seq analysis) were grouped according to predicted functions using EggNOG (v5.0.0). Black bars indicate the percentage of upregulated genes, while gray bars represent the percentage of downregulated genes in cultures grown with *E. citroniae* extract

**Figure 8.**
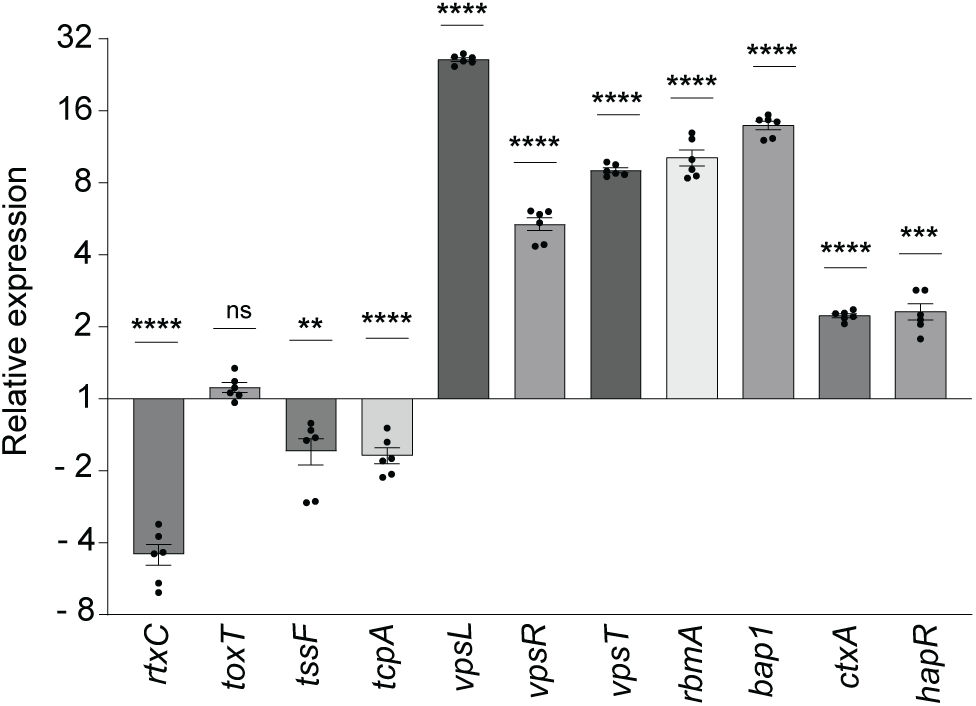
Validation of *V. cholerae* transcriptional responses to *E. citroniae* extracts by quantitative PCR. Relative mRNA levels of selected genes were quantified in *V. cholerae* C6706 grown containing either an extract from *E. citroniae* WAL17108 or a control extract. Expression was normalized to the housekeeping gene *recA* and presented as log₂ fold change relative to the control. Bars represent mean ± SEM from three independent biological replicates. Statistical significance was assessed using an unpaired t-test (*p* < 0.05). Each qRT-PCR reaction was performed in duplicate.

**TABLE 1.** *V. cholerae* genes involved in biofilm formation, virulence, and other relevant pathways that were differentially expressed in response to small molecules produced by *E. citroniae*.

| Annotation | Gene | Fold change |
| --- | --- | --- |
| ***Upregulated*** |  |  |
| periplasmic nitrate reductase subunit alpha | *napA* | 72.0 |
| chaperone NapD |  | 61.8 |
| nitrate reductase cytochrome c-type subunit |  | 52.0 |
| ferredoxin-type protein NapF | *napF* | 51.6 |
| NapC/NirT family cytochrome c |  | 28.9 |
| hypothetical protein |  | 18.9 |
| exopolysaccharide biosynthesis glycosyltransferase VpsL | *vpsL* | 13.4 |
| UDP-N-acetyl-D-mannosamine dehydrogenase VpsB | *vpsB* | 9.56 |
| hypothetical protein | *rbmA* | 9.43 |
| UDP-N-acetylglucosamine 2-epimerase (non-hydrolyzing) VpsA | *vpsA* | 8.97 |
| exopolysaccharide biosynthesis beta-barrel protein VpsM | *vpsM* | 8.70 |
| exopolysaccharide export protein VpsN | *vpsN* | 7.83 |
| beta-prism lectin domain-containing protein | *bap1* | 7.13 |
| exopolysaccharide biosynthesis acetyltransferase VpsC | *vpsC* | 6.37 |
| beta-prism lectin domain-containing protein | *rbmC* | 6.05 |
| exopolysaccharide regulatory protein-tyrosine phosphatase VpsU | *vpsU* | 5.58 |
| O-antigen ligase | *rbmD* | 4.05 |
| exopolysaccharide biosynthesis acetyltransferase VpsG | *vpsG* | 3.71 |
| exopolysaccharide biosynthesis glycosyltransferase VpsD | *vpsD* | 3.68 |
| LuxR family transcriptional regulator VpsT | *vpsT* | 3.38 |
| exopolysaccharide regulatory tyrosine autokinase VpsO | *vpsO* | 3.28 |
| exopolysaccharide biosynthesis protein VpsH | *vpsH* | 3.11 |
| exopolysaccharide biosynthesis glycosyltransferase VpsK | *vpsK* | 3.05 |
| exopolysaccharide biosynthesis flippase VpsE | *vpsE* | 3.00 |
| exopolysaccharide biosynthesis glycosyltransferase VpsI | *vpsI* | 2.80 |
| glycosyl hydrolase family 28-related protein | *rbmB* | 2.78 |
| exopolysaccharide biosynthesis protein VpsJ | *vpsJ* | 2.57 |
| cyclic-di-GMP-binding transcriptional regulator VpsR | *vpsR* | 2.46 |
| exopolysaccharide biosynthesis protein VpsF | *vpsF* | 2.37 |
| ***Downregulated*** |  |  |
| toxin-coregulated pilus major pilin TcpA | *tcpA* | -7.18 |
| RTX toxin-activating lysine-acyltransferase RtxC | *rtxC* | -6.12 |
| toxin-coregulated pilus methyl-accepting chemotaxis protein TcpI | *tcpI* | -5.79 |
| toxin-coregulated pilus pre-pilin peptidase TcpJ | *tcpJ* | -5.60 |
| MARTX multifunctional-autoprocessing repeats-in-toxin holotoxin RtxA | *rtxA* | -4.31 |
| pilus/toxin transcriptional regulator ToxT | *toxT* | -3.69 |
| type VI secretion system baseplate subunit TssF | *tssF* | -3.32 |
| type VI secretion system baseplate subunit TssG | *tssG* | -3.23 |
| toxin-coregulated pilus minor pilin TcpB | *tcpB* | -2.57 |
| type VI secretion system contractile sheath small subunit | *tssB* | -2.51 |
| type VI secretion system contractile sheath large subunit | *tssC* | -2.46 |
| RTX toxin T1SS ABC transporter subunit RtxB | *rtxB* | -2.10 |

### Whole fecal extracts and *E. citroniae* culture extracts display distinct effects on the *V. cholerae* transcriptome

To evaluate how *E. citroniae* metabolites influence *V. cholerae* transcription compared to the effects of the more chemically complex human fecal matrix, we compared our RNA-seq data with our previously published transcriptomic dataset of *V. cholerae* grown in the presence of a human fecal extract (8). A scatter plot comparing the log_2_ fold changes under both conditions is shown in **Figure 9**. This analysis revealed a subset of genes with shared regulation across both conditions, as well as many other genes that respond specifically to *E. citroniae* metabolites. A subset of genes (e.g., the *vps* cluster) that were strongly upregulated in the *E. citroniae* extract remained unchanged or were mildly downregulated in the fecal extract, suggesting specific regulation of biofilm pathways by *E. citroniae* signals. Together, these data indicate that while both fecal and *E. citroniae* specific metabolites markedly modulate *V. cholerae* global transcription, *E. citroniae* metabolites elicits a distinct and more targeted regulatory response compared to the complex and diverse signals present in the whole fecal extract.

**Figure 9.**
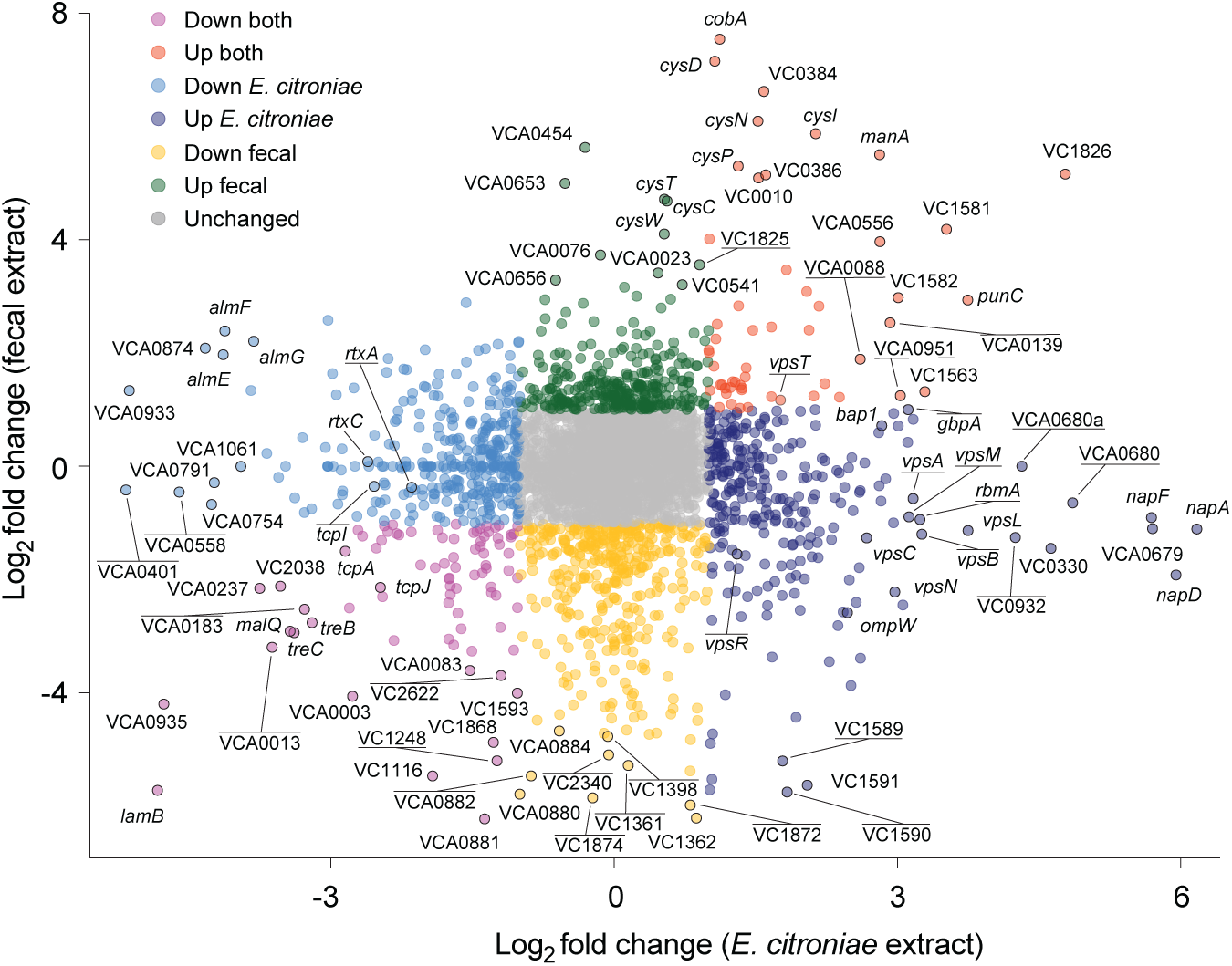
Correlation between transcriptional profiles of V. cholerae grown in the presence of fecal or *E. citroniae* extracts. Each point represents a gene, plotted by its log₂ fold change in *V. cholerae* exposed to *E. citroniae* extract (x-axis) versus its fold change in fecal extract (y-axis). Genes are color-coded by regulation pattern: purple (downregulated in both conditions), orange (upregulated in both), light blue (downregulated in *E. citroniae* extract only), dark blue (upregulated in *E. citroniae* extract only), green (upregulated in fecal extract only), yellow (downregulated in fecal extract only), and gray (not significantly changed in either condition).

### *Vibrio* polysaccharide (*vps*) gene expression is activated by *E. citroniae* culture extracts

To confirm the effect of metabolites produced by *E. citroniae* on *vps* gene expression, we created a reporter system by fusing the promoter region of *vpsR* to the *lux* operon in pBBRLux (12), creating pLC010. The *vpsR* gene encodes a cyclic-di-GMP-binding transcriptional regulator that is critical for the expression of other *vps* genes involved in the synthesis of *Vibrio* polysaccharide, known to be important for biofilm formation (22, 23). In our RNA-seq data, *vpsR* expression was induced 2.46-fold when *V. cholerae* was cultured in the presence of an *E. citroniae* extract. Therefore, we grew our *V. cholerae vpsR* reporter in the presence of a control extract or an *E. citroniae* extract (WAL17108), and measured growth and light production. As can be seen in **Figure 10**, light production was significantly increased when *V. cholerae* was grown in the presence of an *E. citroniae* extract, confirming that *vps* gene expression is activated by compounds produced by *E. citroniae*.

**Figure 10.**
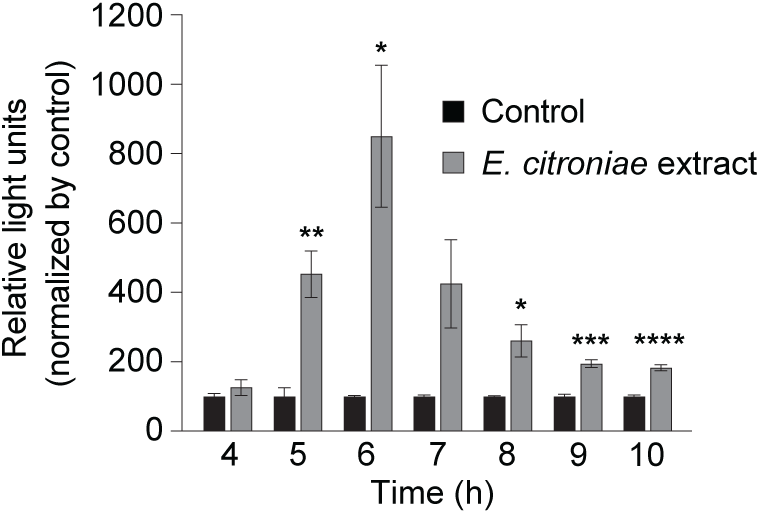
Expression of *vpsR* in a *V. cholerae* reporter strain in response to *E. citroniae* extracts. Light production of by a reporter strain containing a P*vpsR-lux* fusion in the absence and presence of *E. citroniae* extracts (2x concentration). Statistical significance was assessed using Welch’s t test comparing each treatment to the corresponding untreated control. *p < 0.05, **p < 0.005, ***p < 0.0005, ****p < 0.0001

### *E. citroniae* extracts modulate *V. cholerae* interactions with host cells

Due to the observed effects of *E. citroniae* extracts on the expression of several traits that are important for host interactions, we set out to determine if interactions of *V. cholerae* with host cells would be affected by *E. citroniae* culture extracts. Before doing so, we sought to determine if the compounds produced by *E. citroniae* affect host cells, by performing toxicity and viability assays using a human colon epithelial cell line (HT-29). After 24 hours of exposure of HT-29 cells to *E. citroniae* extracts (WAL17108), a concentration-dependent cytotoxic effect was observed, as shown by MTT and LDH assays. MTT results **(Figure 11A**) revealed a significant reduction in cellular metabolic activity when cells were incubated in the presence of *E. citroniae* extracts, as compared to the DMEM control, with a decrease to approximately 5% of the control at the highest concentrations used (10x and 5x), 35% with 2x, and 45% with 1x, remaining significantly lower than the control at all tested concentrations. Interestingly, however, a control extract produced from bacterial culture medium also caused a significant, although less pronounced, reduction in metabolic activity, suggesting that medium components are at least partially responsible for this effect. Nevertheless, it is important to note that a significant difference between the *E. citroniae* and control extracts was still observed at a 2x concentration, the concentration used for most experiments described herein. Similarly, the LDH assay indicated increased membrane damage at higher *E. citroniae* extract concentrations when compared to the DMEM control **(Figure 11B**), with LDH release reaching approximately 50% at the 10x concentration, and decreasing progressively at 5x, 2x, and 1x. In this assay, however, control extracts did not induce any toxicity. Of note, no significant differences were observed between the *E. citroniae* and control extracts at a 2x concentration, the concentration used for most experiments described herein. These findings confirm that, although the effect is somewhat moderate, *E. citroniae* extracts induce dose-dependent cytotoxicity toward HT-29 cells.

**Figure 11.**
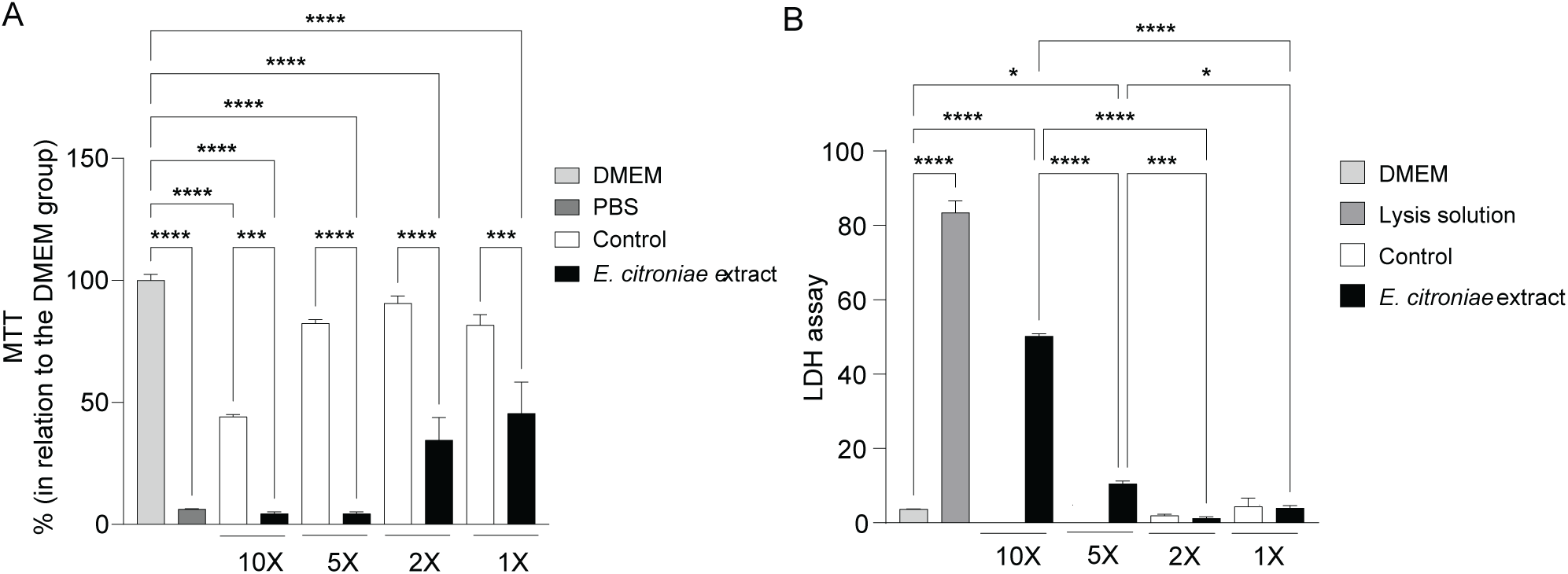
Cytotoxic activity of *E. citroniae* extracts in HT-29 cells. (A) Cell viability was assessed using the MTT assay after treatment with *E. citroniae* extract at concentrations of 10x, 5x, 2x, and 1x. Data are presented as mean ± SEM from five replicates. Statistical significance was determined by one-way ANOVA (*p* < 0.05; **** *p* < 0.001; **** *p* < 0.0001). (B) Cytotoxicity was evaluated using an LDH release assay in HT-29 cells exposed to *E. citroniae* extract at concentrations of 10x, 5x, 2x, and 1x, and expressed as percent cytotoxicity. LDH release from lysed cells served as positive control.

During infection, *V. cholerae* must interact with host cells in the intestinal tract. To determine if bioactive molecules produced by *E. citroniae* modulate these interactions, we performed cell adhesion and invasion assays using *V. cholerae* cells that had been grown in the presence of a control extract or *E. citroniae* extracts (WAL17108). To do this, *V. cholerae* was grown under both conditions until mid-log growth was achieved. Cells were then washed and used to infect HT-29 cells. This was done so the epithelial cells were not exposed to *E. citroniae* extracts, avoiding the issue of toxicity. As can be seen in **Figure 12**, there was a modest, yet noticeable reduction in bacterial counts in both the adhesion and invasion assays in the presence of *E. citroniae* extracts compared to the control. In our adhesion assays, the control group exhibited approximately 2.1×10^5^ CFU/mL, whereas treatment with the *E. citroniae* extracts reduced adhesion to approximately 1.2×10^5^ CFU/mL, representing a significant decrease (**Figure 12A**). Similarly, invasion was significantly reduced in the presence of *E. citroniae* extracts, with CFU counts decreasing from approximately 1.4×10^3^ CFU/mL in the control condition to approximately 8×10^2^ CFU/mL in the group treated with *E. citroniae* extracts (**Figure 11B**). These results indicate that *E. citroniae* extracts inhibit both adhesion and invasion of HT-29 cells by *V. cholerae*. Lastly, in order to evaluate the impact of *E. citroniae* extracts on *V. cholerae* survival following host cell invasion, an overnight *V. cholerae* culture was subcultured in the presence of control or *E. citroniae* extracts until reaching the mid-log growth phase (OD_600_ of approximately 0.5). This time, however, both HT-29 cells and *V. cholerae* were exposed to either condition (control extracts or *E. citroniae* extracts) at the time of infection. However, exposure occurred for only 6 hours to minimize potential cytotoxic effects. At this time point, a significant decrease in *V. cholerae* intracellular counts was still observed in the presence of *E. citroniae* extracts (**Figure 12C**). Also, to ascertain that this result was not a consequence of differences in the numbers of epithelial cells in both conditions, we normalized the number of bacterial CFU to the number of HT-29 cells. This still resulted in significantly reduced numbers of intracellular *V. cholerae* in the presence of *E. citroniae* extracts (**Figure 12D**).

**Figure 12.**
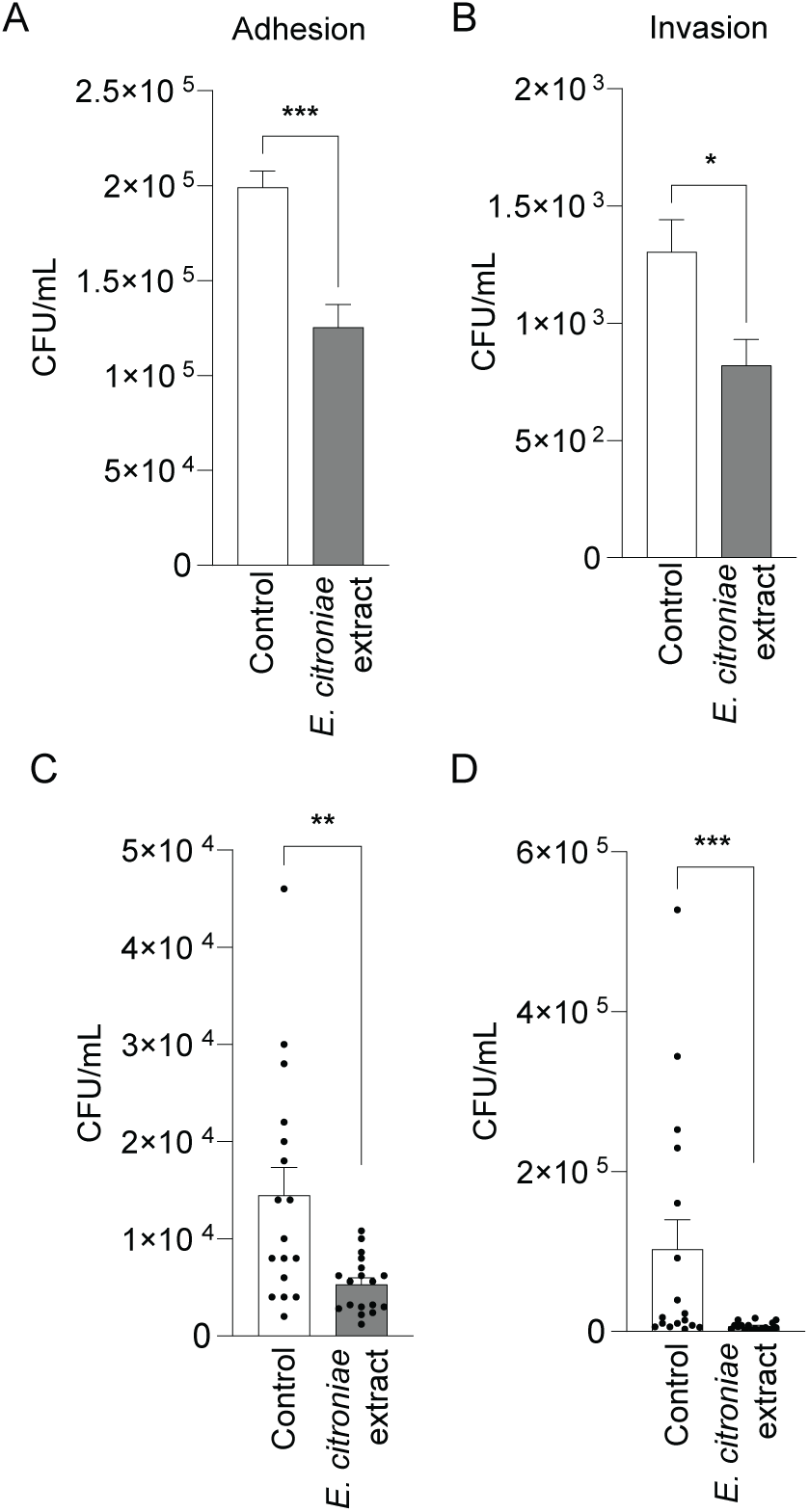
Effect of *E. citroniae* extracts on the adhesion, invasion, and survival of *V. cholerae* in epithelial cells. (A) Representative *V. cholerae* colonies recovered on LB agar plates from the adhesion assay (10⁻² dilution) and invasion assay (10⁰ dilution). (B) Adhesion and invasion of *V. cholerae* to HT-29 cells were quantified after bacteria were grown in the presence or absence of *E. citroniae* extracts (2× concentration). (C) Intracellular survival of *V. cholerae* was assessed 6 hours post-infection. For these assays, bacteria were grown with *E. citroniae* extracts prior to infection and were also exposed to the extracts throughout the infection period. Statistical significance was assessed using unpaired t-tests comparing each treatment to the corresponding untreated control (*p* < 0.05).

## Discussion

Our study demonstrates that metabolites produced by *E. citroniae* profoundly influence the physiology of *V. cholerae*, revealing the role of metabolites produced by the gut microbiota in shaping pathogen behavior. We show that exposure to *E. citroniae* extracts triggers a coordinated shift from a motile, virulent phenotype toward a sessile, biofilm state. This transition is supported by phenotypic assays and transcriptomic data, which together indicate that microbiota-derived metabolites can reprogram *V. cholerae* behavior in ways that may impact colonization and persistence within the host. One of the most striking findings was the strong induction of biofilm formation in the presence of *E. citroniae* extracts, observed under both static and flow conditions. RNA-seq analysis revealed upregulation of the entire *vps* gene cluster, as well as biofilm matrix components such as *rbmA* and *bap1*. These genes are essential for *Vibrio* polysaccharide (VPS) synthesis and biofilm maturation, processes that enhance environmental survival and host colonization (22–24). The activation of *vpsR* and *vpsT*, key regulators of cyclic-di-GMP signaling, further supports the notion that *E. citroniae* metabolites promote a high-c-di-GMP state, favoring biofilm development over motility. This is consistent with our motility assays, which showed a marked reduction in swimming behavior, and aligns with previous reports that microbiota signals can modulate virulence factors in enteric pathogens (2, 7, 8).

Interestingly, besides genes involved in biofilm formation, genes encoding nitrate reductases were among the most highly upregulated in response to *E. citroniae* metabolites, despite the use of aerobic growth conditions. Nitrate respiration is typically associated with anaerobic environments such as the gut lumen and has been linked to enhanced colonization in murine models (25, 26). The activation of this pathway under oxygen replete conditions may suggest that specific metabolites produced by this gut commensal may mimic gut-associated cues, priming *V. cholerae* for survival in niches where oxygen is limited. This metabolic reprogramming, coupled with increased biofilm formation, may represent an adaptive strategy that facilitates long-term persistence within the host or environmental reservoirs.

Besides the aforementioned transcriptional changes, *E. citroniae* metabolites strongly repressed genes that encode major virulence determinants, including the toxin-coregulated pilus (*tcpA*), RTX toxin (*rtxA*, *rtxC*), and components of the type VI secretion system (T6SS). These systems are critical for host colonization, cytotoxicity, and interbacterial competition (27–30). Downregulation of ToxT, the master regulator of virulence gene expression, suggests that *E. citroniae* metabolites interfere with upstream regulatory cascades, potentially reducing the pathogen’s ability to cause acute disease. This dual effect, biofilm promotion and virulence suppression, may reflect a trade-off between transmission and persistence, favoring a less aggressive state under microbiota influence. Conversely, it is possible that the effect of cues produced by *E. citroniae* on *V. cholerae* virulence is more nuanced, and may be involved in regulating the timing, rather than the intensity of the response. Similar antagonistic regulation of motility and virulence by gut metabolites has been reported for other pathogens, but our findings highlight a specific and potent effect of *E. citroniae* metabolites on *V. cholerae* physiology (2, 7, 8).

During *V. cholerae* host colonization and pathogenesis, two early and essential steps are the ability of the bacteria to actively traverse the intestinal mucus layer and adhere to the epithelial cell surface. Our analysis of the *E. citroniae* extracts demonstrated that these metabolites significantly impaired not only swimming motility but also the ability of *V. cholerae* to adhere to epithelial cells, indicating a direct inhibitory effect on another key virulence-associated behavior. Although *V. cholerae* is not considered an invasive organism, our data confirmed earlier reports by showing that *V. cholerae* can invade epithelial cells *in vitro* (31). Interestingly, the ability to invade host cells was also repressed by compounds produced by *E. citroniae*, contributing to the multifaceted nature of the effects elicited by this commensal.

Altogether, the observations presented herein demonstrate the complexity of microbiota-pathogen interactions during cholera infection. While previous studies have shown that certain commensals inhibit *V. cholerae* growth or virulence through short-chain fatty acids or quorum sensing signals (2, 3), our data indicate that *E. citroniae* employs distinct metabolites to modulate biofilm formation and metabolic pathways. Importantly, the ability of microbiota-derived molecules to suppress virulence while promoting biofilm formation could influence disease outcomes by reducing acute pathogenicity yet enhancing environmental persistence, potentially affecting transmission dynamics. These findings underscore the role of microbiota molecules as environmental cues that reprogram *V. cholerae* toward persistence rather than acute infection, potentially reshaping disease dynamics and transmission. By promoting biofilm formation and metabolic adaptations while suppressing virulence, these metabolites may influence both host-pathogen interactions and environmental survival. However, further studies are essential to identify the specific compounds responsible and elucidate their mechanisms of action. Such insights will deepen our understanding of the microbiota-pathogen interplay and pave the way for microbiome-based strategies to mitigate cholera and related enteric infections.

## Acknowledgments

We thank Chris Waters for the kind gift of *V. cholerae* C6706, pBBRLux, and pCMW5. We thank Neta Sal-Man for the kind gift of *V. cholerae* 1276. We also thank Carolina Tropini for kindly sharing *Enterocloster bolteae* strains BAA-613 and CC43_001B, as well as *Enterocloster clostridioformis* strains 2_1_49FAA and WAL7855. We are also grateful to Sarah Vancuren and Jerry He for assistance with growing, preparing, and shipping gut microbiome isolates between laboratories. Research reported in this publication was supported by the National Institute of General Medical Sciences (NIGMS) of the National Institutes of Health under award number P20GM113117. This work used computational resources provided by the RPT04A Bioinformatics Core Facility at Fiocruz, Rio de Janeiro.

---

**Supplementary Figure 1.**
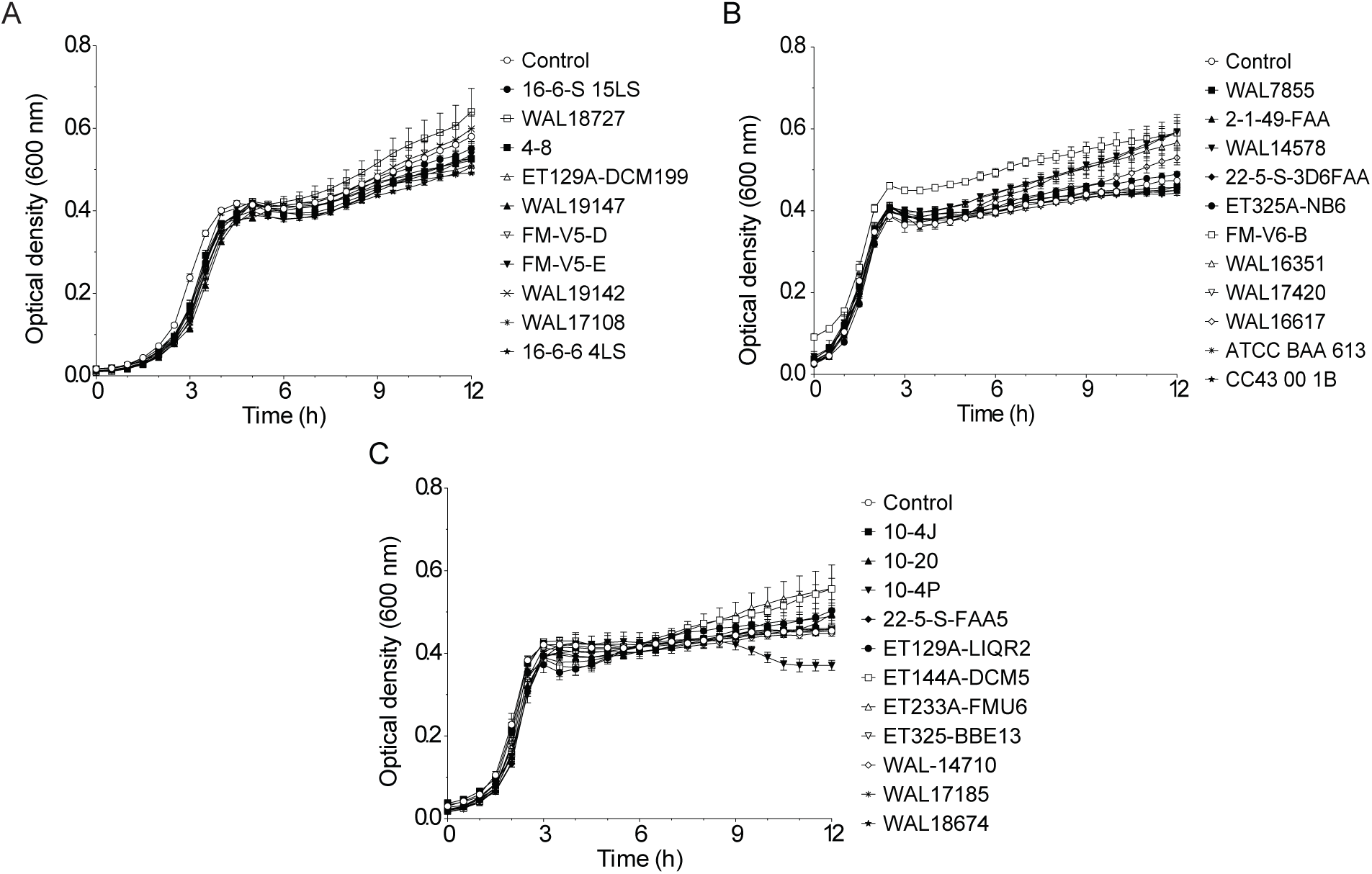
Effect of gut commensal extracts on *V. cholerae* growth. Panels A, B, and C show the results of growth curves performed independently, with different strains of *Enterocloster* spp. and *Clostridium innocuum*.

**Table S1:**
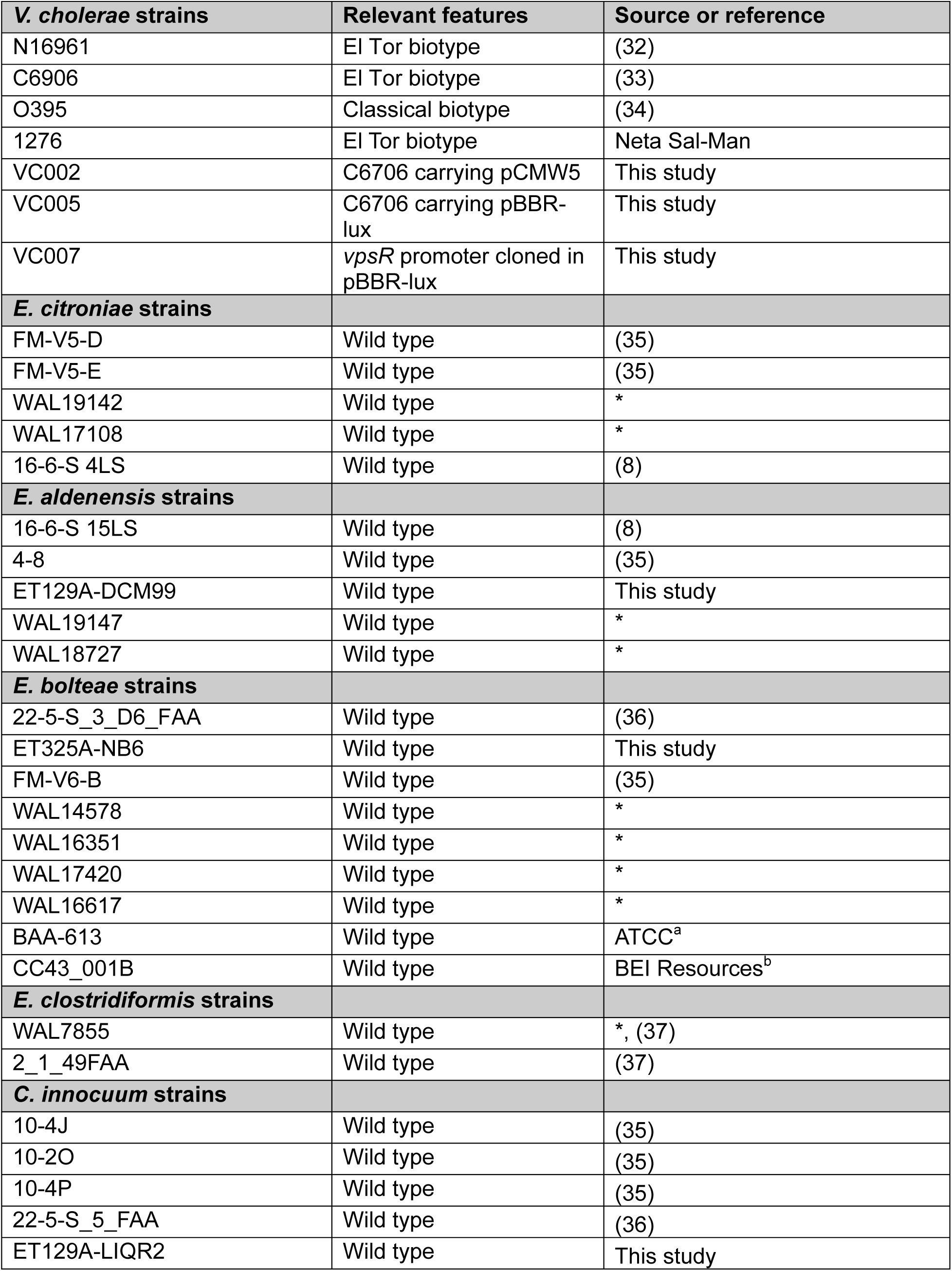
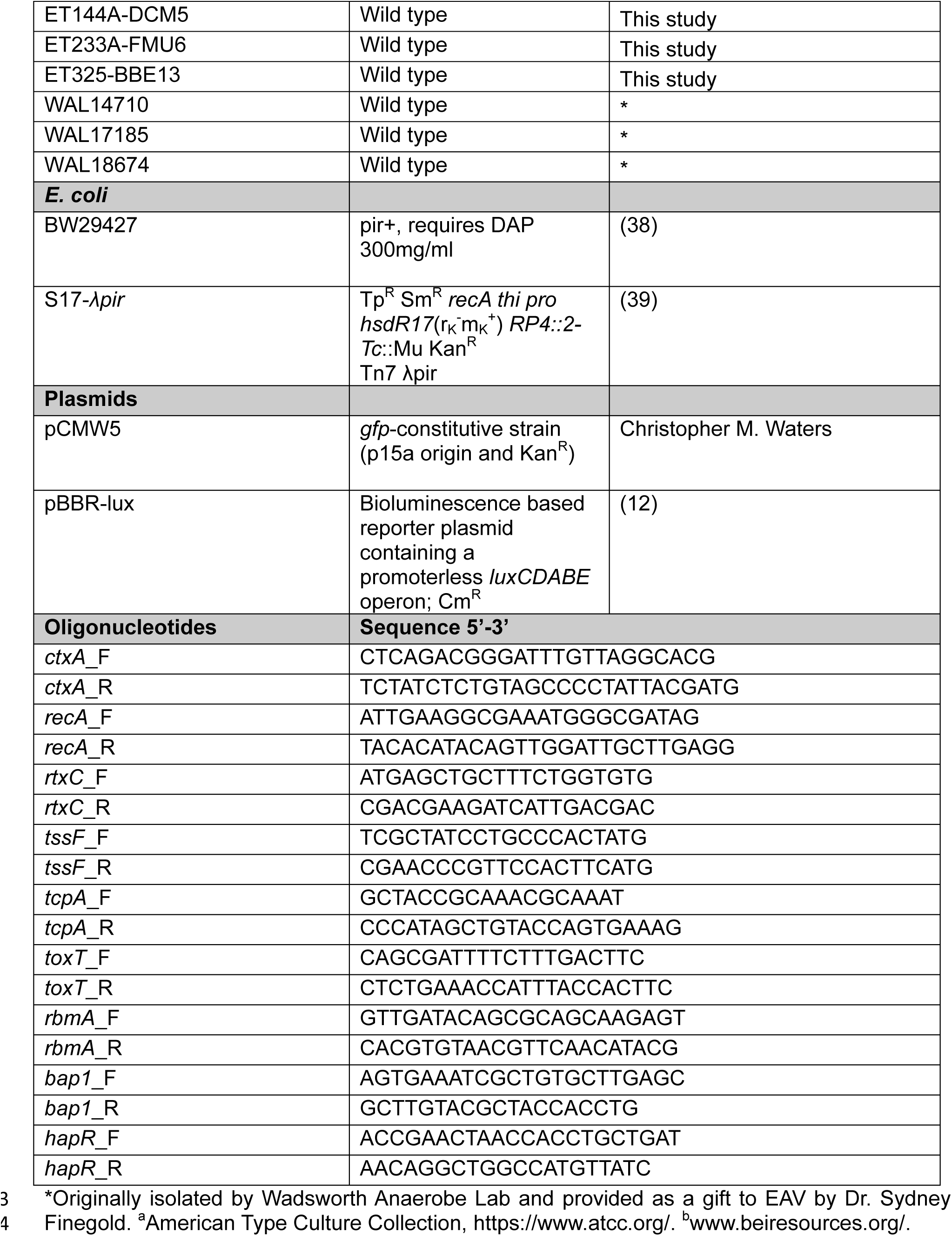
Bacterial strains, plasmids, and oligonucleotides used in this study.

## References

1. WHO. Cholera. Fact sheet: www.who.int/news-room/fact-sheets/detail/cholera. Accessed in November 2025.

2. Hsiao A, Ahmed AM, Subramanian S, Griffin NW, Drewry LL, Petri WA, Jr., et al. Members of the human gut microbiota involved in recovery from *Vibrio cholerae* infection. Nature. 2014;515(7527):423–6.

3. You JS, Yong JH, Kim GH, Moon S, Nam KT, Ryu JH, et al. Commensal-derived metabolites govern *Vibrio cholerae* pathogenesis in host intestine. Microbiome. 2019;7(1):132.

4. Alavi S, Mitchell JD, Cho JY, Liu R, Macbeth JC, Hsiao A. Interpersonal gut microbiome variation drives susceptibility and resistance to cholera infection. Cell. 2020;181(7):1533–46 e13.

5. Zhao W, Caro F, Robins W, Mekalanos JJ. Antagonism toward the intestinal microbiota and its effect on *Vibrio cholerae* virulence. Science. 2018;359(6372):210–3.

6. Yoon MY, Min KB, Lee KM, Yoon Y, Kim Y, Oh YT, et al. A single gene of a commensal microbe affects host susceptibility to enteric infection. Nat Commun. 2016;7:11606.

7. Antunes LC, McDonald JA, Schroeter K, Carlucci C, Ferreira RB, Wang M, et al. Antivirulence activity of the human gut metabolome. mBio. 2014;5(4):e01183–14.

8. Pauer H, Teixeira FL, Robinson AV, Parente TE, De Melo MAF, Lobo LA, et al. Bioactive small molecules produced by the human gut microbiome modulate *Vibrio cholerae* sessile and planktonic lifestyles. Gut Microbes. 2021;13(1):1–19.

9. Peixoto RJM, Alves ES, Wang M, Ferreira RBR, Granato A, Han J, et al. Repression of *Salmonella* host cell invasion by aromatic small molecules from the human fecal metabolome. Appl Environ Microbiol. 2017;83(19).

10. Barrasso K, Chac D, Debela MD, Geigel C, Steenhaut A, Rivera Seda A, et al. Impact of a human gut microbe on *Vibrio cholerae* host colonization through biofilm enhancement. Elife. 2022;11.

11. Campbell IW, Dehinwal R, Morano AA, Dailey KG, Zingl FG, Waldor MK. *Vibrio cholerae* motility is associated with inter-animal transmission. Nat Commun. 2025;16(1):7989.

12. Hammer BK, Bassler BL. Regulatory small RNAs circumvent the conventional quorum sensing pathway in pandemic *Vibrio cholerae*. Proc Natl Acad Sci USA. 2007;104(27):11145–9.

13. O’Toole GA. Microtiter dish biofilm formation assay. J Vis Exp. 2011(47).

14. Tinevez JY, Perry N, Schindelin J, Hoopes GM, Reynolds GD, Laplantine E, et al. TrackMate: An open and extensible platform for single-particle tracking. Methods. 2017;115:80–90.

15. Ewels P, Magnusson M, Lundin S, Kaller M. MultiQC: summarize analysis results for multiple tools and samples in a single report. Bioinformatics. 2016;32(19):3047–8.

16. Dobin A, Davis CA, Schlesinger F, Drenkow J, Zaleski C, Jha S, et al. STAR: ultrafast universal RNA-seq aligner. Bioinformatics. 2013;29(1):15–21.

17. Love MI, Huber W, Anders S. Moderated estimation of fold change and dispersion for RNA-seq data with DESeq2. Genome Biol. 2014;15(12):550.

18. Livak KJ, Schmittgen TD. Analysis of relative gene expression data using real-time quantitative PCR and the 2-DDCt method. Methods. 2001;25(4):402–8.

19. Marin MA, Fonseca EL, Andrade BN, Cabral AC, Vicente AC. Worldwide occurrence of integrative conjugative element encoding multidrug resistance determinants in epidemic *Vibrio cholerae* O1. PLoS One. 2014;9(9):e108728.

20. Rekha K, Venkidasamy B, Samynathan R, Nagella P, Rebezov M, Khayrullin M, et al. Short-chain fatty acid: an updated review on signaling, metabolism, and therapeutic effects. Crit Rev Food Sci Nutr. 2024;64(9):2461–89.

21. Huerta-Cepas J, Szklarczyk D, Heller D, Hernandez-Plaza A, Forslund SK, Cook H, et al. eggNOG 5.0: a hierarchical, functionally and phylogenetically annotated orthology resource based on 5090 organisms and 2502 viruses. Nucleic Acids Res. 2019;47(D1):D309–D14.

22. Fong JCN, Syed KA, Klose KE, Yildiz FH. Role of *Vibrio* polysaccharide (*vps*) genes in VPS production, biofilm formation and *Vibrio cholerae* pathogenesis. Microbiology. 2010;156(Pt 9):2757–69.

23. Yildiz FH, Dolganov NA, Schoolnik GK. VpsR, a member of the response regulators of the two-component regulatory systems, is required for expression of *vps* biosynthesis genes and EPS(ETr)-associated phenotypes in *Vibrio cholerae* O1 El Tor. J Bacteriol. 2001;183(5):1716–26.

24. Hsieh ML, Waters CM, Hinton DM. VpsR directly activates transcription of multiple biofilm genes in *Vibrio cholerae*. J Bacteriol. 2020;202(18).

25. Heidelberg JF, Eisen JA, Nelson WC, Clayton RA, Gwinn ML, Dodson RJ, et al. DNA sequence of both chromosomes of the cholera pathogen *Vibrio cholerae*. Nature. 2000;406(6795):477–83.

26. Bueno E, Sit B, Waldor MK, Cava F. Anaerobic nitrate reduction divergently governs population expansion of the enteropathogen *Vibrio cholerae*. Nat Microbiol. 2018;3(12):1346–53.

27. Schuhmacher DA, Klose KE. Environmental signals modulate ToxT-dependent virulence factor expression in *Vibrio cholerae*. J Bacteriol. 1999;181(5):1508–14.

28. Linhartova I, Bumba L, Masin J, Basler M, Osicka R, Kamanova J, et al. RTX proteins: a highly diverse family secreted by a common mechanism. FEMS Microbiol Rev. 2010;34(6):1076–112.

29. Fu Y, Waldor MK, Mekalanos JJ. Tn-Seq analysis of *Vibrio cholerae* intestinal colonization reveals a role for T6SS-mediated antibacterial activity in the host. Cell Host Microbe. 2013;14(6):652–63.

30. Fast D, Kostiuk B, Foley E, Pukatzki S. Commensal pathogen competition impacts host viability. Proc Natl Acad Sci USA. 2018;115(27):7099–104.

31. Panigrahi P, Tall BD, Russell RG, Detolla LJ, Morris JG, Jr. Development of an *in vitro* model for study of non-O1 *Vibrio cholerae* virulence using Caco-2 cells. Infect Immun. 1990;58(10):3415–24.

32. Levine MM, Black, R.E., Clements, M.L., Nalin, D.R., Cisneros, L., Finkelstein, R.A. Volunteer studies in development of vaccines against cholera and enterotoxigenic *Escherichia coli*: a review. In: T. Holme JH, M. H. Merson, R. Mollby, editor. Acute Enteric Infections in Children: New Prospects for Treatment and Prevention. Amsterdam: Elservier; 1981. p. 449–59.

33. Weng Y, Bina XR, Bina JE. Complete genome sequence of *Vibrio cholerae* O1 El Tor strain C6706. Microbiol Resour Announc. 2021;10(3).

34. Mekalanos JJ. Duplication and amplification of toxin genes in *Vibrio cholerae*. Cell. 1983;35(1):253–63.

35. Oliphant K, Parreira VR, Cochrane K, Allen-Vercoe E. Drivers of human gut microbial community assembly: coadaptation, determinism and stochasticity. ISME J. 2019;13(12):3080–92.

36. Yen S, McDonald JA, Schroeter K, Oliphant K, Sokolenko S, Blondeel EJ, et al. Metabolomic analysis of human fecal microbiota: a comparison of feces-derived communities and defined mixed communities. J Proteome Res. 2015;14(3):1472–82.

37. Human Microbiome Jumpstart Reference Strains C, Nelson KE, Weinstock GM, Highlander SK, Worley KC, Creasy HH, et al. A catalog of reference genomes from the human microbiome. Science. 2010;328(5981):994–9.

38. Marzluf GA, Reddy CA, Snyder LR, Beveridge TJ, Breznak JA, Schmidt TM. Methods for general and molecular microbiology. 3rd ed. Washington: ASM Press 2007.

39. de Lorenzo V, Timmis KN. Analysis and construction of stable phenotypes in Gram-negative bacteria with Tn5- and Tn10-derived minitransposons. Methods Enzymol. 1994;235:386–405.

